# Distinct neurogenic progenitor cell populations balance cell type production in the embryonic mouse retina

**DOI:** 10.64898/2026.08.03.738451

**Authors:** Henry L. Bushnell, Constance L. Cepko

## Abstract

Mammalian vision depends on the reliable production of over 100 neural cell types from multipotent retinal progenitor cells. How so many cell types are generated in the correct proportions throughout the retina remains incompletely understood. Focusing on the early embryonic period in mice, we found that retinal neurogenesis is organized through discrete neurogenic progenitor cell (NPC) populations with distinct fate biases. NPCs expressing *Galanin* primarily produce retinal ganglion cells and amacrine cells, whereas *Olig2*-expressing NPCs produce cones and horizontal cells in the same temporal window. Clonal analysis revealed that these two fate-biased NPC populations arise predominantly from asymmetric, renewing progenitor cell divisions that produce one fate-biased NPC daughter cell. Differential Notch signaling regulates the production of these NPC populations, with *Galanin*^+^ and *Olig2*^+^ NPCs representing Notch^High^ and Notch^Low^ states, respectively. Manipulating Notch signaling was sufficient to toggle their production. The expression of Notch signaling components across NPCs is consistent with a lateral inhibition mechanism, in which fate-biased NPCs promote neighboring cells to adopt a complementary fate-biased NPC state. These findings suggest that the robust production of diverse retinal cell types is achieved through local feedback that balances discrete, fate-biased NPC populations.

## INTRODUCTION

A central question in developmental biology is how cellular diversity is reliably generated during development. In some systems, invariant lineages and stereotyped divisions yield highly predictable outcomes (Lemaire, 2009; Sulston et al., 1983). In mammals, however, many tissues arise through more probabilistic progenitor behaviors, in which individual lineage trajectories vary substantially (Chan et al., 2019; Pijuan-Sala et al., 2018). How such variable developmental programs nevertheless produce every cell type in correct proportions remains unresolved.

The mammalian retina provides a powerful model for this problem. It contains more than 100 neuronal cell types generated from a common pool of multipotent retinal progenitor cells (RPCs) during embryogenesis (Turner et al., 1990). Because tangential migration is limited (Reese et al., 1995; Turner et al., 1990), these cell types must be specified locally and in appropriate proportions across the tissue to support visual function. Decades of work have identified transcription factors required for individual retinal cell fates (Clark et al., 2019; Elliott et al., 2008; Ge et al., 2023; Xiang, 2013), revealing the importance of gene regulation in constraining or enabling specific cell identities. By contrast, much less is known about how individual cells select among multiple fate options, or how these decisions are coordinated locally to ensure robust tissue-level cell type composition.

Here, we examine cell fate choice during the early embryonic window in mouse (E12–E16), when RPCs concurrently generate retinal ganglion cells (RGCs), amacrine cells, horizontal cells, and cone photoreceptors (Cepko, 2014; Young, 1985). Clonal studies have shown that single RPCs can produce highly variable combinations of cell classes (Turner et al., 1990), yet retinal development remains strikingly reproducible. Whether this robustness emerges through communication among progenitor cells or through largely cell-intrinsic mechanisms remains a key unresolved question.

Fate-biased progenitor states have been described in the developing retinas of several species (Brzezinski et al., 2011; Brzezinski et al., 2012; Hafler et al., 2012; Emerson et al., 2013; Schick et al., 2019; Chen and Emerson, 2021; Wang et al., 2020; Nerli et al., 2023). In mouse, an *Olig2*-expressing progenitor population generates cones and horizontal cells during early embryogenesis through terminal neurogenic divisions (Hafler et al., 2012). Related cone/horizontal-biased progenitor cells have also been identified in chick through activity of the ThrbCRM1 enhancer (Emerson et al., 2013; Schick et al., 2019), suggesting that retinal neurogenesis may proceed through discrete intermediate progenitor states with biased output. Yet analogous progenitor populations that preferentially generate RGCs or amacrine cells have not been defined in mammals. This gap is notable because these classes comprise a major fraction of retinal cell type diversity, including more than 40 molecularly distinct RGC subtypes identified by single-cell transcriptomics (Goetz et al., 2022). How cells of this diverse cell class are reliably produced remains unclear.

Notch signaling is a strong candidate regulator of fate decisions in the early embryonic retina. Across developing tissues, Notch coordinates neighboring progenitor cell behaviors (Gozlan and Sprinzak, 2023; Louvi and Artavanis-Tsakonas, 2006). In the retina, Notch has been implicated in controlling progenitor cell maintenance, proliferation, and fate choice (Henrique et al., 1997; Jadhav et al., 2006a; Jadhav et al., 2006b; Mills and Goldman, 2017; Mizeracka et al., 2013; Nelson et al., 2007; Yaron et al., 2006). Canonically, Notch ligand-receptor interactions between adjacent cells trigger release of the Notch receptor intracellular domain (NICD), which enters the nucleus to activate transcriptional programs. In the embryonic retina, most insight has come from loss-of-function studies, where reduced Notch activity promotes differentiation and frequently favors cone production (Chen and Emerson, 2021; Jadhav et al., 2006b; Kaufman et al., 2019; Luo et al., 2012; Yaron et al., 2006). However, perturbation of distinct pathway components yields variable effects on other early-born cell classes, particularly RGCs, pointing to a more complex role in fate choice (Austin et al., 1995; Bosze et al., 2020; Bosze et al., 2023; Lee et al., 2005; Riesenberg and Brown, 2016; Riesenberg et al., 2009; Takatsuka et al., 2004; Zheng et al., 2009).

Here, we seek to address how progenitor cells produce diverse cell types during early embryonic retinogenesis. We define transcriptional heterogeneity among neurogenic progenitor cells (NPCs) in the early embryonic mouse retina and identify a *Galanin* (*Gal*)-expressing NPC population distinct from previously described *Olig2*^+^ NPCs. *Gal*^+^ NPCs undergo terminal divisions and are biased to generate RGCs and amacrine cells. Rather than emerging as sibling cells in the same division, *Gal*^+^ and *Olig2*^+^ NPCs arise in asymmetric divisions that generate an RPC sibling. These neurogenic populations correspond to Notch^High^ and Notch^Low^ signaling states, respectively, and express

Notch pathway components in patterns consistent with a lateral inhibition mechanism. Manipulating Notch activity was sufficient to shift the specification of these NPC populations, suggesting that neighboring progenitor cells balance local cell type production by inducing adjacent cells to adopt cell states with complementary fate biases.

## RESULTS

### Expression of *Gal* and *Olig2* mark distinct progenitor populations in the embryonic retina

In the early embryonic retina, progenitor cells generate post-mitotic neurons of four distinct cell classes. While a fate-biased NPC population that produces two of these classes (cones and horizontal cells) has been identified by the expression of *Olig2* (Hafler et al., 2012), the progenitor of RGCs and amacrine cells born in this temporal window remains undefined (Figure 1A).

**Figure 1.**
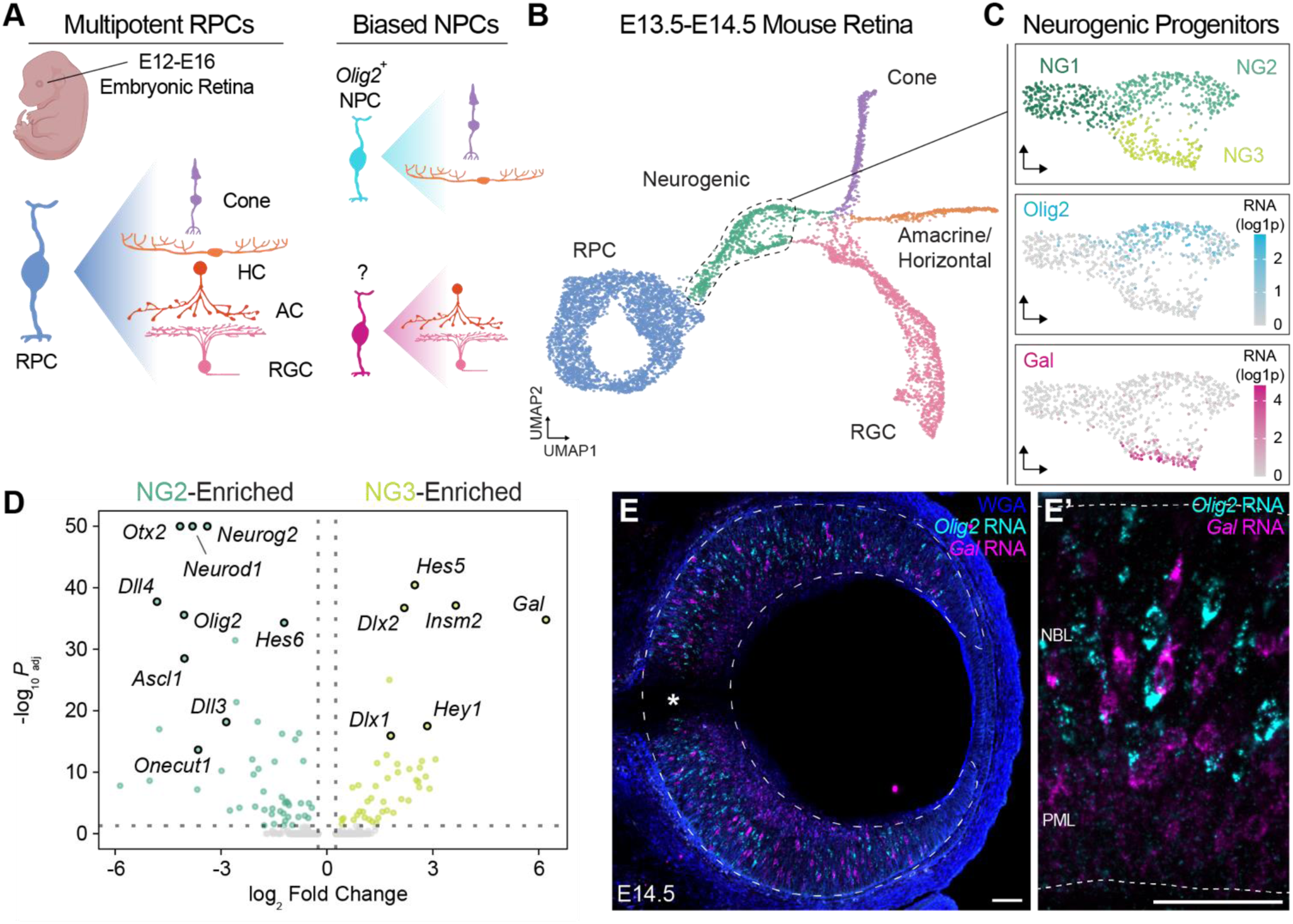
***Gal* and *Olig2* expression mark distinct progenitor populations in the embryonic retina** (A) Schematic illustrating RPC competence during early embryonic development. A subset of progenitor cells, which express *Olig2*, exhibit restricted competence, whereas additional progenitor states remain uncharacterized. (B) UMAP of integrated scRNA-seq datasets from Wu et al. 2021 and Balasubramanian et al. 2021. (C) UMAP of potential NPCs colored by cluster identity (top), *Olig2* expression (middle), and *Gal* expression (bottom). (D) Volcano plot showing differentially expressed genes between clusters NG2 and NG3. (E) E14.5 retinal section with HCR RNA-FISH for *Olig2* (cyan) and *Gal* (magenta) with WGA (blue) marking cell boundaries. Scale bar, 50 µm. Dotted line and asterisk indicate retinal boundary and optic nerve, respectively. (E’) Higher-magnification view of the E14.5 retina showing the neuroblast layer (NBL) and post-mitotic layer (PML). Dashed lines indicate the retinal boundaries.

To investigate the heterogeneity of NPCs in this period, we integrated published scRNA-seq datasets from E13.5 and E14.5 mouse retinas (Balasubramanian et al., 2021; Wu et al., 2021) (Figure 1B; 7,016 cells total). Louvain clustering resolved all major embryonic cell classes (Figure 1B), with balanced integration across both datasets (Supp. Figure 1A). We identified a cluster of NPCs, defined by their co-expression of proliferation marker *Top2a* and NPC marker *Atoh7* (Wu et al., 2021), as well as the expression of neurogenic transcription factors *Ascl1*, *Neurog2*, *Neurod2*, and *Olig2* (Supp. Figure 1B-D).

Further Louvain clustering of the NPC population revealed three distinct subpopulations: an S-phase cluster (NG1), an *Olig2*-expressing cluster (NG2), and a third cluster expressing G2/M marker genes but lacking *Olig2* expression (NG3) (Figure 1C). We hypothesized that NG2 and NG3 may correspond to distinct NPC populations that are poised to produce different cell classes. Consistent with this hypothesis, NG2 cells had increased expression of *Olig2* (log_2_FC = 4.05, p_adj_ = 2.6 x 10^-36^), a known marker of cone/horizontal-biased NPCs, as compared to NG3. Furthermore, transcription factors known to promote the cone and horizontal cell fates (e.g. *Otx2*, *Neurod1*, *Onecut1*) were highly expressed by NG2 versus NG3 cells (Figure 1D). NG3 cluster cells were marked by expression of RGC fate determinants *Dlx1* (Zhang et al., 2017), *Dlx2*, and *Isl1* (Figure 1D), although both NG2 and NG3 cells expressed other RGC and amacrine cell transcriptional effectors such as *Atoh7*, *Sox4*, and *Six3* (Supp. Figure 2). Interestingly, the strongest marker of NG3 cells was *Galanin* (*Gal*; log_2_FC = 6.21, p_adj_ = 1.6 x 10^-35^), a neuropeptide known to be expressed during retinogenesis (Brodie-Kommit et al., 2021; Weir et al., 2021). While Gal is dispensable for RGC genesis (Brodie-Kommit et al., 2021), we hypothesized that we could use Gal as a genetic handle to access this specific population of progenitor cells.

**Figure 2.**
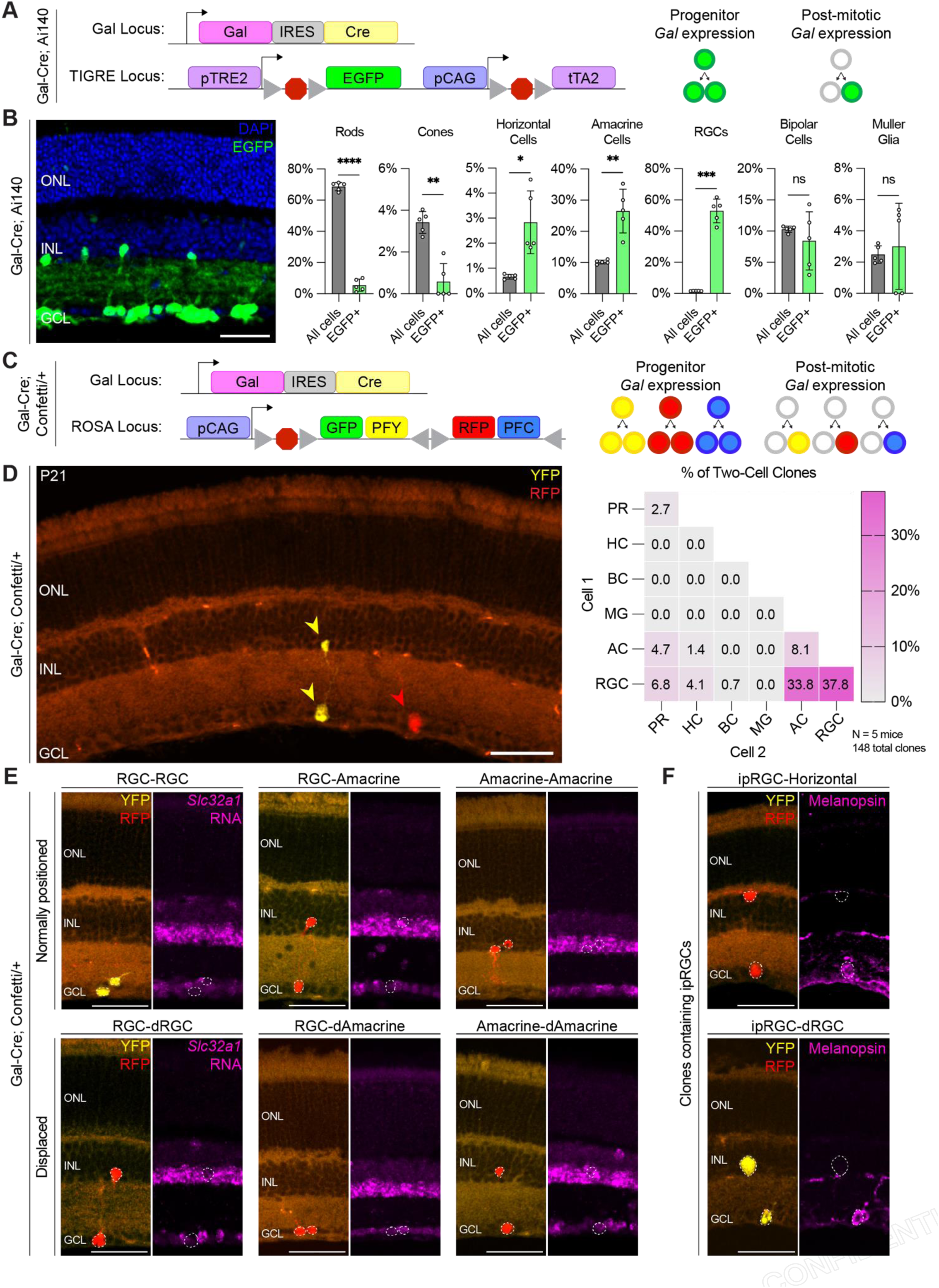
***Gal^+^* NPCs generate RGCs and amacrine cells** (A) Schematic of alleles in Gal-Cre; Ai140 mice. (B) (Left) P21 retinal section from a Gal-Cre; Ai140 mouse stained with DAPI (blue). Scale bar, 50 µm. (Right) Quantification of retinal cell types in the total population (gray) and among EGFP^+^ cells (green). Each dot represents one mouse (n = 5); for each mouse, the plotted value is the mean of five fields of view within a single retina. Data are mean ± SD. Paired, two-tailed t tests were used to assess differences between the total and EGFP^+^ fractions across matched mice. ns, not significant; *p<0.05, **p<0.01, ***p<0.001, ****p<0.0001. (C) Schematic of alleles in Gal-Cre; Confetti/+ mice. (D) (Left) P21 retinal section from a Gal-Cre; Confetti/+ mouse showing YFP (yellow) and RFP (red). Scale bar, 50 µm. Arrows indicate cells with productive Confetti reporter recombination. (Right) Quantification of cell types among two-cell clones, shown as the percentage of total two-cell clones (N = 5 mice, 148 total clones). (E) Example two-cell clones of the three most frequent clone compositions from Gal-Cre; Confetti/+ mice with normally positioned cells (Top) and clones containing a displaced cell (Bottom). Images show YFP (yellow) and RFP (red) with HCR RNA-FISH for amacrine cell marker *Slc32a1* (magenta). Scale bars, 50 µm. (F) Example two-cell clones from Gal-Cre; Confetti/+ mice with intrinsically photosensitive RGCs (ipRGCs). Images show YFP (yellow) and RFP (red) with anti-Melanopsin immunostaining (magenta). Scale bars, 50 µm.

To validate that *Gal*-expressing NPCs constitute a distinct population of progenitor cells *in vivo*, we used Hybridization Chain Reaction RNA fluorescence *in situ* hybridization (HCR RNA-FISH). We detected *Gal*^+^ cells in the central retina as early as E12.5, coinciding with the onset of neurogenesis (Supp. Figure 3). Many *Gal*^+^ cells were productively labeled within 30 minutes of EdU administration, indicating that they are in the S/G2-phase of the cell cycle. By E14.5, *Gal* and *Olig2* transcripts were detected throughout the neuroblast layer, and were confirmed to be expressed by mutually exclusive cell populations (Figure 1E). Low *Gal* expression was also seen in the post-mitotic layer, consistent with previous observations that *Gal* is expressed by nascent RGCs (Brodie-Kommit et al., 2021; Clark et al., 2019). These data suggest that *Gal* expression marks a unique NPC population that is potentially specialized to generate a cellular repertoire distinct from that of *Olig2*^+^ NPCs.

**Figure 3.**
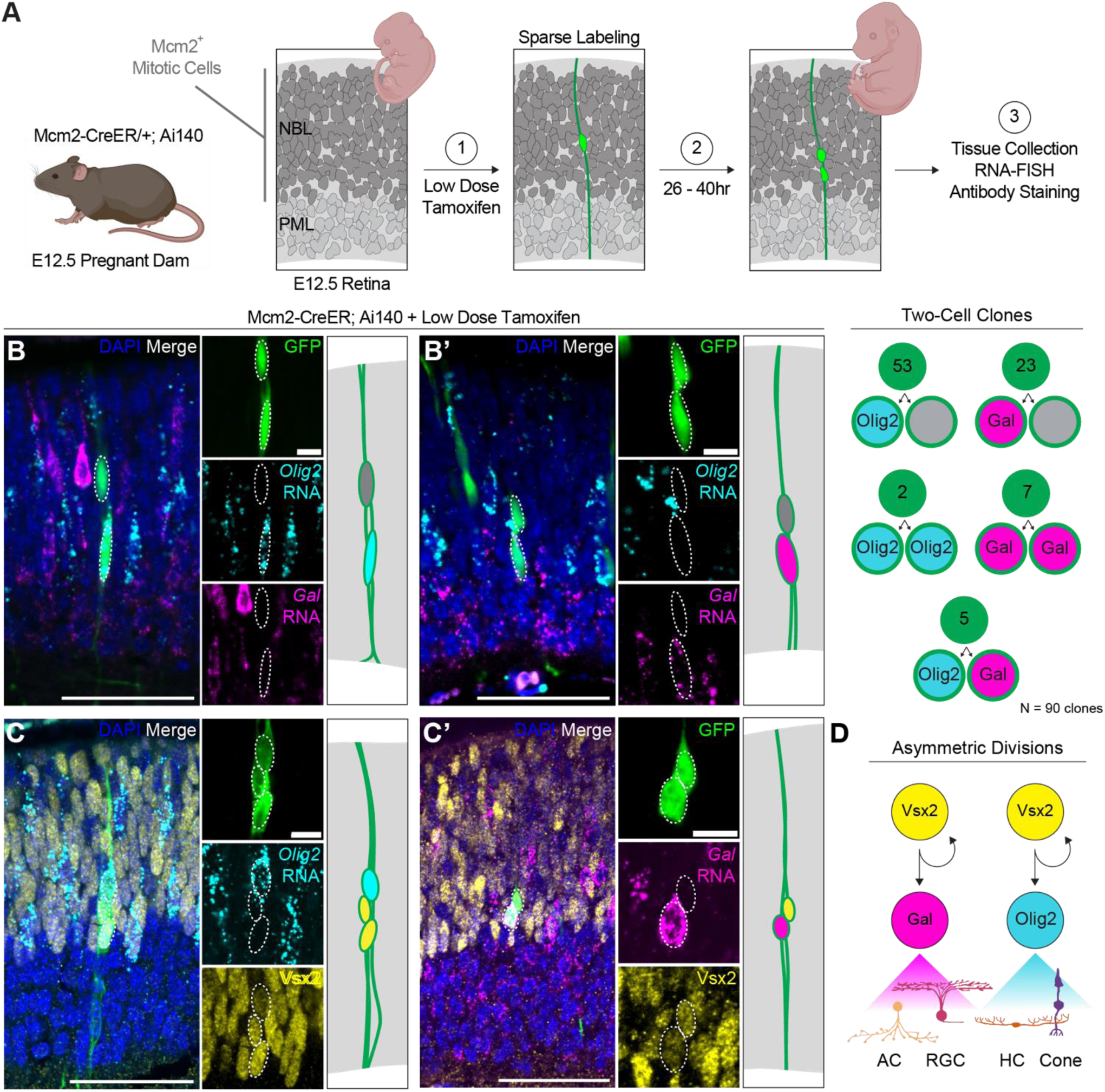
***Gal*^+^ and *Olig2*^+^ NPCs arise from asymmetric divisions with an RPC sibling** (A) Schematic of the strategy used to generate sparsely labeled clones in the embryonic mouse retina. (B, B’) Representative two-cell clones from Mcm2-CreER; Ai140 retinal sections following low-dose tamoxifen administration, containing an *Olig2*^+^ cell (B) or a *Gal*^+^ cell (B’). Sections were analyzed by HCR RNA-FISH for *Olig2* (cyan) or *Gal* (magenta) with DAPI (blue) marking cell nuclei. Scale bars, 50µm (low magnification) and 10 µm (high magnification). (Right) Summary schematics of the two-cell clone compositions observed (n = 90 two-cell clones). (C, C’) Representative clones containing an *Olig2*^+^ cell (C) or a *Gal*^+^ cell (C’), analyzed by HCR RNA-FISH for *Olig2* (cyan) or *Gal* (magenta) together with immunostaining for Vsx2 (yellow) and DAPI staining (blue). Scale bars, 50µm (low magnification) and 10 µm (high magnification). (D) Model of fate-biased progenitor cell production by asymmetric division.

### *Gal^+^* NPCs terminally divide to produce RGCs and amacrine cells

After identifying a population of NPCs transcriptionally distinct from the cone/horizontal-biased *Olig2*^+^ NPCs, we sought to determine if *Gal*^+^ cells have a similar fate bias. We employed a Cre-based labeling strategy to determine which cell types are generated from *Gal^+^* NPCs. A mouse strain with Cre integrated into the Gal locus (Gal-Cre) was crossed with a Cre reporter mouse strain (Ai140) to permanently label *Gal*-expressing cells and their progeny with Enhanced Green Fluorescent Protein (EGFP) (Figure 2A, B). Retinas from Gal-Cre; Ai140 mice were collected at P21, after retinal development is complete. All major retinal cell classes were identified by cell body localization and HCR RNA-FISH for marker gene expression (Supp. Figure 4). While the overall proportion of each cell type in Gal-Cre; Ai140 retinas was consistent with wild-type proportions (Jeon et al., 1998), EGFP-labeled cells were significantly enriched for RGCs, amacrine cells, and horizontal cells (Figure 2B). In contrast, both rod and cone photoreceptors were significantly underrepresented, suggesting that most photoreceptors are not generated from a *Gal*-expressing progenitor cell. Although bipolar cells and Müller glia were marked at rates similar to their proportions in the retina, these cell classes are generated postnatally(Young, 1985) and thus their marking does not reflect production from embryonic *Gal*^+^ NPCs.

**Figure 4.**
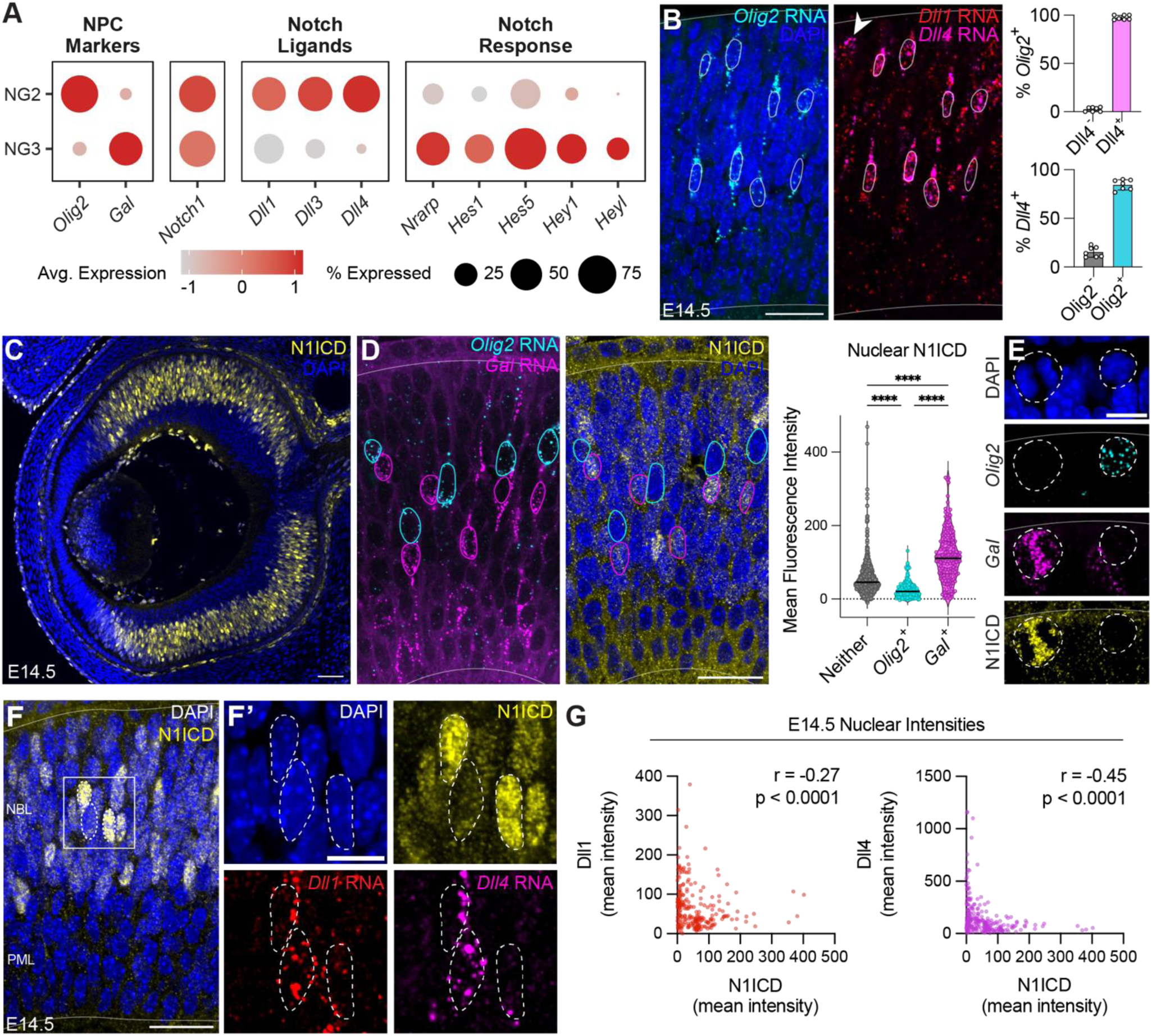
Differential Notch signaling activity in *Gal*^+^ and *Olig2*^+^ NPCs. (A) Dot plot of Notch receptor, ligand, and target gene expression across NPC subclusters. (B) (Left) E14.5 retinal section stained with DAPI (blue) and analyzed by HCR RNA-FISH for *Olig2* (cyan), *Dll1* (red), and *Dll4* (magenta). Circles indicate the nuclear boundary of *Olig2*^+^ cells; solid lines indicate the retinal boundary. Scale bar, 10 µm. White arrow indicates Dll4^+^Olig2^-^ cell. (Right) Quantification of *Dll4* positivity in *Olig2*^+^ cells and *Olig2* positivity in *Dll4*^+^ cells. n = 7 retinal sections. Bars are mean ± SD. (C) E14.5 retinal section immunostained for cleaved N1ICD (yellow) with DAPI (blue). Scale bar, 50 µm. (D) Representative E14.5 retinal section analyzed by HCR RNA-FISH for *Gal* (magenta) and *Olig2* (cyan) together with immunostaining for cleaved N1ICD (yellow). Dotted lines indicate nuclear boundaries for *Gal*^+^ and *Olig2*^+^ cells. Scale bar, 50µm. (Right) Quantification of nuclear cleaved N1ICD fluorescence intensity in NBL cells, stratified by marker expression. Data are shown as violin plots with individual values overlaid (each dot represents one cell); black bars indicate the mean. Statistical significance was determined by Kruskal-Wallis test followed by Dunn’s multiple comparisons test; **** *p* < 0.0001. (E) Example of mitotic *Gal*^+^ (magenta) and *Olig2*^+^ (cyan) cells with DAPI (blue) and immunostaining for cleaved N1ICD (yellow). Scale bar, 10 µm. (F, F’) E14.5 retinal section immunostained for cleaved NICD (yellow) with DAPI (blue) (F) and corresponding high-magnification view of the same section showing HCR RNA-FISH for *Dll1* (red) and *Dll4* (magenta) (F’). Dotted lines indicate the nuclear boundaries of cells of interest. Scale bars, 50 µm (low magnification) and 10 µm (high magnification). (G) Scatter plots of mean nuclear fluorescence intensity from individual high-expressing cells in the E14.5 retina (n = 248 cells from 3 retinas), filtered to include only cells in the top 10% of N1ICD, Dll1, or Dll4 signal intensity. Spearman correlation coefficients and p-values for N1ICD vs. Dll1 (Left) or N1ICD vs. Dll4 (Right) are indicated on each plot.

To distinguish whether *Gal* expression history reflects reporter recombination in mitotic cells versus post-mitotic cells, we crossed Gal-Cre mice to the Confetti reporter (Gal-Cre; Confetti), which expresses one of four fluorophores upon Cre-mediated *loxP* recombination (Figure 2C). Consistent with the reported low productive recombination efficiency of the Confetti reporter with weak Cre drivers, P21 retinas from Gal-Cre; Confetti mice had infrequent, well-separated labeled cells (Figure 2D). Labeled cells were predominantly RFP^+^ or YFP^+^ with rare CFP^+^ and GFP^+^ cells, in agreement with reported recombination bias of the Confetti reporter (Snippert et al., 2010). Given the observed sparse labeling (∼18 labeled cells per 30µm full retinal section) and limited tangential dispersion of retinal clones (Reese et al., 1995; Turner et al., 1990), same-color cells that were radially aligned (separated by ≤ 10µm) were identified as putative sibling cells derived from a single *Gal*^+^ progenitor cell.

Labeled cells were mostly observed as singlets (∼90%), which may reflect post-mitotic *Gal* expression, developmental apoptosis of labeled cells, or a delay between Gal locus activation and Cre expression and subsequent reporter recombination. The remaining ∼10% of recombination events were radial pairs of labeled cells. Same-color groupings larger than two cells were rare (<1%, Supp. Figure 5). This suggests that *Gal*^+^ NPCs typically undergo a single, terminal division, like *Olig2*^+^ NPCs(Hafler et al., 2012). Among two-cell clones (n = 5 mice, 148 total clones), the most frequent cell type compositions were RGC-RGC (37.8%), RGC-amacrine (33.8%), and amacrine-amacrine (8.1%), with additional pairings observed at lower frequencies (Figure 2D). Notably, Müller glia were absent from two-cell clones, consistent with the interpretation that their labeling in Gal-Cre; Ai140 retinas was likely due to post-mitotic Gal expression. These clone compositions confirm that *Gal*^+^ NPCs predominantly generate RGCs and amacrine cells through a terminal division.

**Figure 5.**
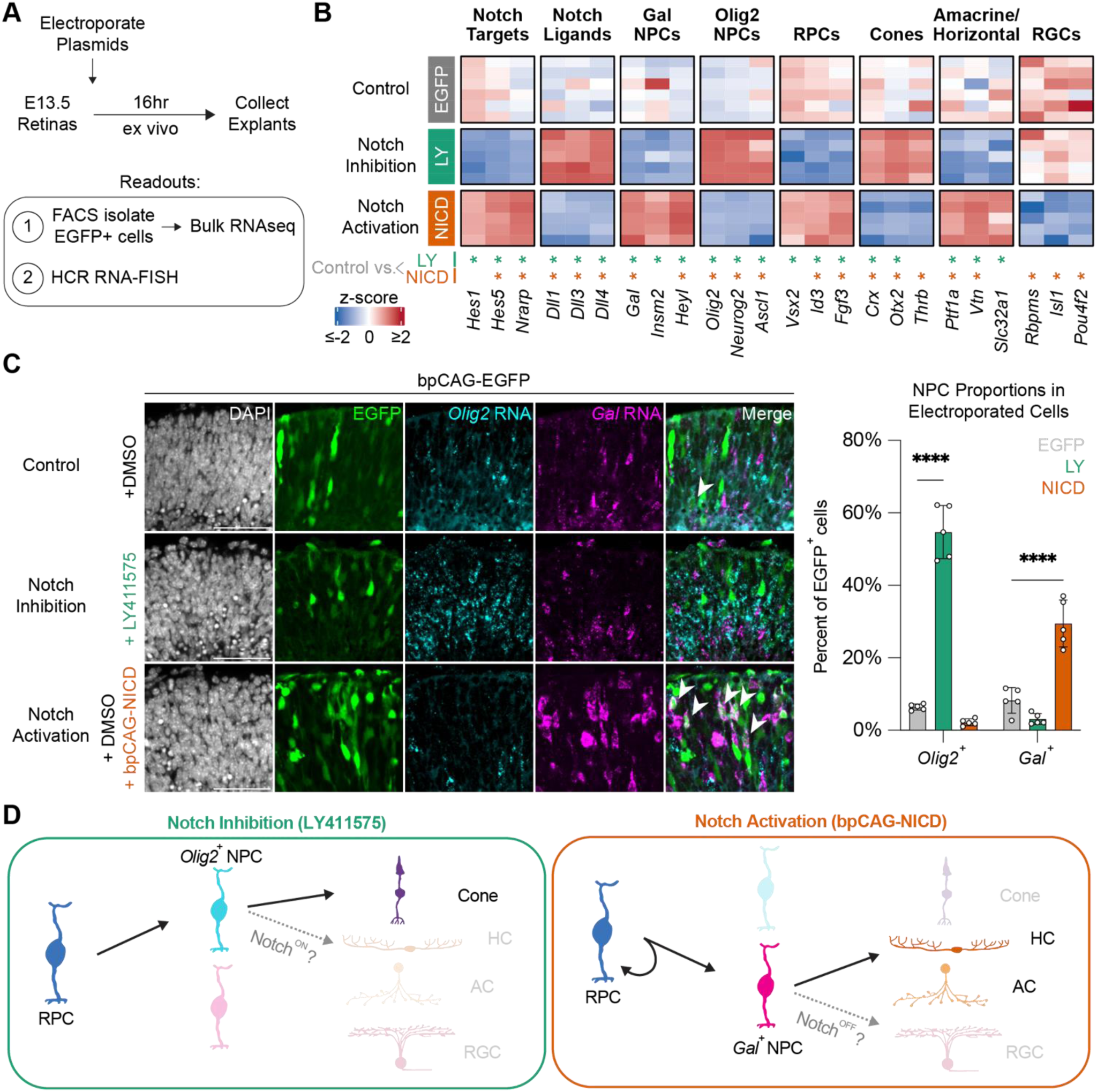
Notch activity shifts production of NPC populations. (A) Schematic of the experimental workflow for Notch signaling manipulation in embryonic retinal explants. (B) Heatmap of gene expression from bulk RNA-seq of FACS-isolated EGFP^+^ cells after 16 hr explant culture. Replicates are shown as rows; color indicates gene-wise z-scored normalized expression. (Bottom) Asterisks denote differential expression at adjusted p<0.05 (DESeq2) for control versus LY (green) and control versus NICD (orange) comparisons. (C) Sections of E13.5 retinal explants after 16 hours in culture with the indicated Notch perturbations. DAPI (white), EGFP (green), and HCR RNA-FISH for *Olig2* (cyan) and *Gal* (magenta) are shown. Arrows indicate examples of EGFP^+^ cells co-expressing *Gal*. Scale bar, 50 µm. (Right) Quantification of the percentage of EGFP^+^ cells expressing *Gal* or *Olig2* across conditions. n = 5 retinas. Bars are mean ± SD. Statistical significance was determined by two-way ANOVA followed by Dunnett’s multiple-comparisons test; ****p < 0.0001. (D) Model summarizing the effects of Notch perturbation on early embryonic retinal cell populations.

A mysterious feature of RGCs and amacrine cells is that a subset of each cell class is spatially “displaced”. The majority of amacrine cells are located in the inner nuclear layer, though a portion are found in the ganglion cell layer, where they account for roughly half of all cells (Choi et al., 2023; Jeon et al., 1998). Likewise, ∼2% of RGCs are displaced to the inner nuclear layer (Dräger and Olsen, 1981; Duda et al., 2025). The developmental origins of these displaced populations remain unclear. To assess whether displaced cells can arise from *Gal*^+^ NPCs, we performed HCR RNA-FISH on Gal-Cre; Confetti retinas for *Slc32a1*, a marker of amacrine cells (Choi et al., 2023), to distinguish cell classes in the ganglion cell layer and inner nuclear layer (Figure 2E). We observed both displaced amacrine cells and displaced RGCs within two-cell clones, indicating that they can be generated by *Gal*^+^ NPCs.

Both types of displaced cells were found in diverse pairings, which suggests that displaced cells are not generated in stereotyped progenitor divisions.

As mentioned above, RGCs are highly diverse. The broad RGC labeling in Gal-Cre; Ai140 retinas suggests that many different RGC subtypes are generated by *Gal*^+^ NPCs. One RGC type of interest is intrinsically photosensitive RGCs (ipRGCs), which express melanopsin (Opn4), an opsin protein more closely related to the rhabdomeric opsins typically found in invertebrate photoreceptors (Guido et al., 2022; Sexton et al., 2012). ipRGCs are essential for circadian photoentrainment and other non-image-forming light responses (Berson et al., 2002; Hattar et al., 2003; Panda et al., 2003). Although regulators of ipRGC differentiation have been described(Mao et al., 2014), the progenitor cells that produce ipRGCs remain uncharacterized. Using melanopsin immunostaining, we identified several ipRGCs within two-cell clones in Gal-Cre; Confetti retinas (Figure 2F), indicating that ipRGCs can arise from *Gal*^+^ NPCs. Sibling cells of ipRGCs were varied but fell within the major outputs of *Gal*^+^ NPCs (RGCs, amacrine cells, and horizontal cells). This suggests that ipRGCs are born from fate-biased, but multipotent, NPCs rather than a dedicated, ipRGC-specific progenitor cell.

### *Gal*^+^ and *Olig2*^+^ NPCs arise from asymmetric divisions with an RPC sibling

The production of different cell types by *Gal*^+^ and *Olig2*^+^ NPCs suggests that some progenitor cells are fate-biased upon entry into distinct NPC states. Patterns of NPC genesis have not been described in mammals, so it is unclear whether these fate biases extend back through multiple cell divisions, i.e., represent sublineages dedicated to the production of different cell types. We therefore asked whether *Gal*^+^ and *Olig2*^+^ NPCs can be generated as sibling cells from a common division, or whether they arise separately through either symmetric divisions (producing two NPCs of the same type) or asymmetric divisions (producing a single NPC alongside a non-NPC sibling). To investigate these clonal relationships, progenitor cells were sparsely labeled by administering a low dose of tamoxifen to E12.5 dams carrying embryos with an Mcm2-CreER allele, which is expressed in mitotic cells (Maslov et al., 2007; West et al., 2022), and the Ai140 Cre reporter. Embryos were collected 26-40 hours after tamoxifen administration to allow reporter recombination and progenitor cell proliferation. Pairs of EGFP^+^ cells were identified as clones if they were aligned within the same radial column and well-spaced from other EGFP^+^ cells. We then used HCR RNA-FISH to identify clones containing at least one *Gal*^+^ or *Olig2*^+^ cell (Figure 3A).

Across 90 such two-cell clones, the sibling of a *Gal*^+^ or *Olig2*^+^ cell most frequently expressed neither *Gal* nor *Olig2* (n = 76/90 clones; Figure 3B). Homotypic pairs (*Gal*^+^-*Gal*^+^ or *Olig2*^+^-*Olig2*^+^) were observed but were infrequent (n = 9/90 clones) and may reflect post-mitotic marker expression after NPC division or incomplete marker transcript degradation in newly post-mitotic cells. Heterotypic *Gal*^+^-*Olig2*^+^ pairs were uncommon (n = 5/90; <6%), indicating that while these two NPC populations can arise as sibling cells, they do so infrequently. Instead, the sibling of a *Gal*^+^ or *Olig2*^+^ cell often expressed high levels of the RPC marker *Vsx2* (Figure 3C), consistent with asymmetric divisions that produce a fate-biased NPC alongside a multipotent RPC sibling (Figure 3D).

Asymmetric RPC division could support a model in which deeper RPC lineages are biased to asymmetrically produce only *Gal*^+^ or *Olig2*^+^ NPCs; however, single embryonic RPC clones often contain cell types associated with the outputs of both fate-biased NPC populations (Turner et al., 1990). Therefore, we propose a model in which multipotent RPCs sequentially produce different types of fate-biased NPCs through successive asymmetric divisions.

### NPC populations display hallmarks of Notch lateral inhibition

The production of *Gal*^+^ and *Olig2*^+^ NPCs through asymmetric RPC divisions must be balanced to ensure the correct proportion and diversity of mature retinal cell types. Whether the proportions of NPC populations are controlled cell-intrinsically or through intercellular coordination is unclear. The local production of divergent cell states often occurs through Notch lateral inhibition, in which neighboring cells reinforce opposing Notch transcriptional states through neighbor-neighbor ligand-receptor interactions (Gozlan and Sprinzak, 2023; Heitzler and Simpson, 1991; Shimojo et al., 2011). In studies of embryonic retinal development, Notch pathway manipulation often results in contradictory outcomes on terminal cell fate choice. We hypothesize that an underexplored and significant role of Notch may be the balancing of neighboring NPC populations, which ultimately determines terminal cell fate distributions.

scRNA-seq data was consistent with a Notch lateral inhibition mechanism in which Notch^Low^ cells express Notch ligands to induce a Notch^High^ state in adjacent cells. Differential expression analysis of the E13.5-E14.5 scRNA-seq dataset revealed broad differences in components of the Notch signaling pathway between *Gal*^+^ and *Olig2*^+^ NPC populations (Figure 1D). Strikingly, *Olig2*^+^ NPCs were enriched for the Delta-family ligands *Dll1*, *Dll3*, and *Dll4* (Figure 4A), and represented the dominant source of Notch ligand expression in the embryonic dataset (Supp. Figure 6). *Olig2*^+^ NPCs specifically express high levels of *Dll4* compared to *Gal*^+^ NPCs (log_2_FC = 4.81, p_adj_ = 1.8 x 10^-38^).

We validated this relationship *in vivo* at E14.5 (97.6 ± 2.3% of *Olig2*^+^ cells expressed *Dll4*, Figure 4B). Furthermore, *Dll4* expression was confirmed to be unique to *Olig2*^+^ cells (84.5 ± 5.4% of *Dll4*^+^ cells co-expressed *Olig2*; n = 7 sections from 2 retinas; Figure 4B). Rare *Dll4*^+^*Olig2*^−^ cells were apically located, consistent with continued *Dll4* expression in newly post-mitotic photoreceptors. Together, these findings identify *Olig2*^+^ NPCs as a major source of Notch ligands in the embryonic retina.

We therefore hypothesized that *Olig2*^+^ NPCs occupy a Notch^Low^ state that activates Notch signaling in neighboring progenitor cells. Consistent with this model, *Gal*^+^ and *Olig2*^+^ NPCs expressed similar levels of Notch1, the principal embryonic retinal receptor, yet canonical Notch targets *Hes1*, *Hes5*, *Hey1*, *Heyl*, and *Nrarp* were selectively enriched in *Gal*^+^ NPCs and depleted in *Olig2*^+^ NPCs (Figure 1D, Figure 4A).

We sought to directly measure Notch activity *in situ*. To distinguish real-time Notch receptor activation from downstream pathway activity (Bosze et al., 2020), we used an antibody specific to the cleaved Notch1 Intracellular Domain (N1ICD), which is generated upon receptor activation by Notch ligands on neighboring cells. N1ICD localized to nuclei and displayed a salt-and-pepper pattern across the neuroblast layer, and was largely absent from the post-mitotic layer (Figure 4C). This pattern indicates sharp local heterogeneity in Notch signaling among neighboring cells. Quantification of N1ICD staining in different populations showed that *Gal*^+^ NPCs had significantly higher nuclear N1ICD than either *Olig2*^+^ NPCs or double-negative cells in the neuroblast layer (Figure 4D). *Olig2*^+^ cells exhibited the lowest N1ICD levels of all groups, demonstrating that, compared to surrounding RPCs, *Gal*^+^ and *Olig2*^+^ NPCs correspond to Notch^High^ and Notch^Low^ states, respectively. Although prior work suggested reduced Notch activity during mitosis (Nelson et al., 2007), mitotic *Gal*^+^ NPCs frequently retained strong N1ICD signal (Figure 4E).

A defining feature of feedback-driven lateral inhibition is reciprocal patterning, whereby Delta-ligand^High^ Notch^Low^ cells neighbor Delta-ligand^Low^ Notch^High^ cells. To test for this pattern directly, we combined detection of *Dll1* and *Dll4* transcripts with N1ICD immunostaining in the same tissue sections. While dense cell processes make it challenging to determine exactly which cells a ligand-expressing cell may be able to signal with, we repeatedly observed adjacent cell bodies with complementary states: *Dll1*/*4*^High^ N1ICD^Low^ cells directly neighboring *Dll1*/*4*^Low^ N1ICD^High^ cells (Figure 4F). Across cells with detectable pathway activity or ligand expression (top 10% of N1ICD, *Dll1*, or *Dll4* signal), N1ICD levels were inversely related to Delta-family ligand abundance (N1ICD vs. *Dll1*: Spearman r = -0.27, P < 0.0001, N1ICD vs. *Dll4*: Spearman r = -0.45, P < 0.0001; Figure 4G). These data support a lateral inhibition mechanism in which Delta-ligand^High^ *Olig2*^+^ NPCs promote a *Gal*^+^ NPC state (Notch^High^), whereas Delta-ligand^Low^ *Gal*^+^ NPCs permit adjacent progenitor cells to adopt an *Olig2*^+^ NPC state (Notch^Low^).

### Notch activity shifts production of NPC populations

Our results suggested that divergent Notch signaling activity was associated with different fate-biased NPC populations. To directly test whether Notch signaling causally specifies NPC populations, we manipulated Notch activity in embryonic retinal explants. Following dissection, E13.5 retinas were electroporated with an EGFP expression plasmid (bpCAG-EGFP), to preferentially label mitotically active cells (Matsuda and Cepko, 2004), and cultured ex vivo for 16 hours (Figure 5A). Notch signaling was inhibited by addition of the γ-secretase inhibitor LY411575, or activated by co-electroporation of an NICD expression plasmid (bpCAG-NICD). EGFP^+^ cells were isolated by FACS and profiled by bulk RNA-seq (n = 5-6 biological replicates per condition).

Given the low levels of nuclear N1ICD in *Olig2*^+^ NPCs *in vivo*, we hypothesized that Notch inhibition would increase the specification of *Olig2*^+^ NPCs. Indeed, Notch inhibition using LY411575 upregulated transcripts characteristic of *Olig2*^+^ NPCs and photoreceptor differentiation. Conversely, NICD expression increased the abundance of transcripts associated with *Gal*^+^ NPCs, RPC proliferation, and nascent amacrine and horizontal cell identity (Figure 5B). Neither perturbation induced RGC markers. Instead, NICD expression significantly reduced levels of *Rbpms*, *Isl1*, and *Pou4f2*, suggesting that while Notch activity can promote the *Gal*^+^ NPC state, the subsequent transition to the RGC fate or expression of RGC differentiation markers may require downregulation of Notch activity. Similarly, although *Olig2*^+^ NPCs generate horizontal cells (Hafler et al., 2012), sustained Notch inhibition reduced expression of early horizontal cell markers, suggesting that horizontal cell differentiation requires dynamic Notch signaling activity rather than prolonged Notch inhibition (Figure 5B).

These results suggest that Notch causally affects transcription associated with NPC states. To assess whether these transcriptional changes reflected a meaningful change in the distribution of fate-biased NPCs, we performed HCR RNA-FISH on electroporated explants. Notch inhibition increased the fraction of EGFP^+^ cells expressing *Olig2* from 6.4 ± 0.6% to 54.7 ± 7.4% (Figure 5C; n = 5 retinas). Conversely, NICD expression increased the *Gal*^+^ fraction from 8.3 ± 3.5% to 29.5 ± 6.4%. In each condition, induction of one NPC marker was accompanied by a reduction in the other. Thus, Notch activity is sufficient to shift specification between fate-biased NPC populations (Figure 5D).

## DISCUSSION

Our findings support a model in which embryonic retinal neurogenesis is organized through distinct, Notch-regulated NPC states. We identified *Gal*-expressing NPCs, which predominantly produce RGCs and amacrine cells, as a population orthogonal to *Olig2*-expressing NPCs, which are biased to generate cones and horizontal cells. Both states arise predominantly through asymmetric progenitor divisions that couple production of a fate-biased NPC with maintenance of an RPC sibling. We showed that Notch activity distinguishes these NPC states, and Notch pathway manipulation was sufficient to shift production between them. Together, these results suggest that retinal neurogenesis is locally diversified by progenitor cell interactions that stabilize complementary fate-biased NPC states.

### A *Galanin*-expressing neurogenic population biased to produce RGCs and amacrine cells

A longstanding question in retinal development is how variable clonal outputs robustly produce the diverse cell types found throughout the mammalian retina. Lineage tracing has shown that RPCs can generate all major retinal classes, yet individual clonal compositions are not stereotyped (Turner et al., 1990). However, analyses of smaller clones suggest a more ordered structure to neurogenic divisions, implying that they are not entirely random. Such intermediate progenitor states are best defined in zebrafish (He et al., 2012; Nerli et al., 2020; Nerli et al., 2023; Wang et al., 2020), and the identification of cone/horizontal-biased NPCs in chick (Emerson et al., 2013) and mouse (Brzezinski et al., 2011; Hafler et al., 2012) suggests that discrete neurogenic intermediates are a conserved feature of vertebrate retinogenesis.

The identification of *Gal*^+^ NPCs reveals previously unresolved heterogeneity within the early embryonic mouse retina. While prior evidence for RGC-biased progenitors in mouse retina was largely indirect or reliant on the non-specific NPC marker *Atoh7* (Brzezinski et al., 2012; Yang et al., 2003), *Gal* expression now provides a prospective marker for this neurogenic population and enables direct clonal interrogation of its outputs. *Gal*^+^ NPC clones frequently contained RGC-RGC or RGC-amacrine pairs, but additional sibling combinations were also observed. These variable outputs indicate that neurogenic bias does not impose a single invariant fate program. For instance, rare two-cell clones containing ipRGCs arose from *Gal*^+^ NPCs while retaining the broader clonal variability characteristic of the *Gal*^+^ NPC state. Together, these findings suggest that neurogenic programs may constrain broad neuronal class identity while terminal subtype specification remains more flexible or is resolved later in development.

Sparse clonal analysis indicated that *Gal*^+^ and *Olig2*^+^ NPCs are typically generated through asymmetric RPC divisions that produce a fate-biased NPC and a multipotent RPC sibling. This architecture provides a simple mechanism by which the embryonic retina may couple tissue expansion to progressive neurogenesis through sequential cell divisions. Distinct NPC states with different biases likely emerge at later stages of retinogenesis. For example, *Olig2*^+^ NPCs present at birth produce rods and amacrine cells rather than cones and horizontal cells (Hafler et al., 2012), illustrating that NPC competency evolves over developmental time. This developmental strategy resembles the well-studied production of *Drosophila* ganglion mother cells, whose stage-specific competencies are directed by a cascade of temporal transcription factors (Brody and Odenwald, 2000; Isshiki et al., 2001). Connecting the shifting competencies of RPCs with the discrete NPC states they produce is an important area for future investigation.

### Notch activity directs distinct NPC populations

We show that Notch signaling directs the specification of distinct NPC populations. Expression patterns between neighboring NPCs are consistent with lateral inhibition, a feedback mechanism in which fate bias in one progenitor cell influences the complementary bias of neighboring cells. We propose this as a mechanism for cell type balancing during retinal neurogenesis.

Notch signaling activity varies among neighboring cells, but within neurogenic populations this variability resolves into two discrete states: *Gal*^+^ NPCs exhibit high Notch activity and *Olig2*^+^ NPCs exhibit low Notch activity. Perturbation experiments demonstrate that this difference is causal, as Notch activation promotes the *Gal*^+^ NPC state while inhibition promotes the *Olig2*^+^ NPC state. Delta-family ligands are enriched in NPCs, with *Olig2*^+^ NPCs representing a major source of ligand expression in the early embryonic retina. Because *Olig2*^+^ NPCs are themselves Notch^Low^, this ligand expression positions them to activate Notch in neighboring progenitor cells in a lateral inhibition mechanism that reinforces opposite transcriptional states between adjacent cells. Reciprocal activation of Notch target genes in *Gal*^+^ NPCs is consistent with this model, in which signaling feedback between fate-biased NPCs actively shapes local neurogenic output (Figure 6A).

**Figure 6.**
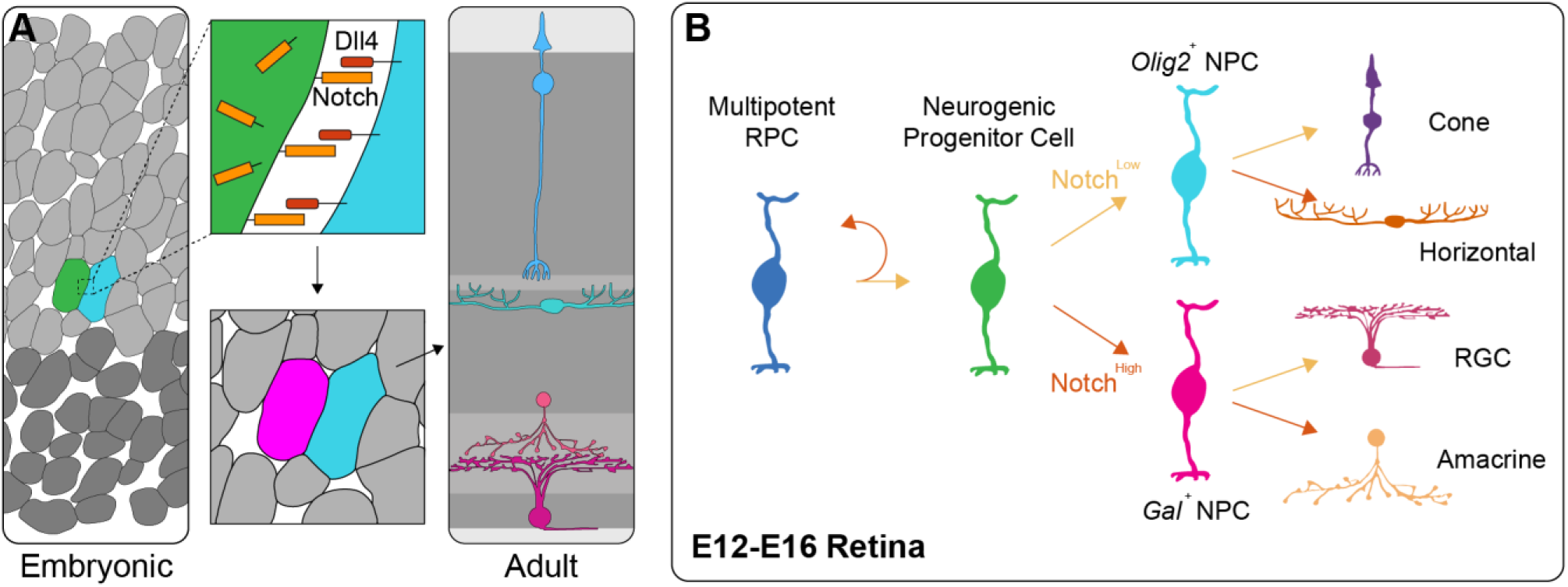
Model of embryonic neurogenesis. (A) Model of lateral inhibition-mediated diversification of neighboring NPC states with complementary competencies. (B) Diagram of potential Notch-mediated branchpoints in embryonic cell fate specification.

These results provide a framework for reconciling the diverse outcomes reported from Notch perturbation studies in the retina. Rather than reflecting inconsistencies in pathway function, these differences likely arise because Notch acts at multiple sequential decision points during neurogenesis. We propose that there may be as many as three sequential Notch^High^/Notch^Low^ branchpoints in embryonic retinal development (Figure 6B). First, the RPC decision of whether to proliferate, which is well established to be Notch-regulated and may involve oscillatory signaling dynamics (Ivanov, 2019; Kageyama et al., 2008; Manning et al., 2019; Shimojo et al., 2008). Second, the choice between RGC/amacrine-biased and cone/horizontal-biased neurogenic programs, which we show here is directed by Notch. Third, we speculate that there may be another Notch-gated decision influencing the progeny of a fate-biased NPC. In this context, daughters of *Olig2*^+^ NPCs may be specified as cones (Notch^Low^) or horizontal cells (Notch^High^), and the daughters of *Gal*^+^ NPCs may be specified as RGCs (Notch^Low^) or amacrine cells (Notch^High^).

This logic of sequential, Notch-gated fate restrictions promotes local cell type diversity through neighboring cell feedback without requiring deterministic lineage programs.

## SUPPLEMENTAL FIGURES

### LEAD CONTACT

Requests for further information and resources should be directed to and will be fulfilled by the lead contact, Connie Cepko.

## MATERIALS AVAILABILITY

Plasmids generated in this study are available upon request. Mouse lines used in this study are available from The Jackson Laboratory (Ai140, R26R-Confetti, Mcm2-CreERT2) and Charles River (CD-1 IGS). Gal-IRES-Cre mice were provided by the laboratory of Dr. Heike Münzberg-Gruening and were obtained under an MTA (availability subject to MTA terms).

## DATA AND CODE AVAILABILITY

This paper analyzes existing, publicly available data, accessible at the Gene Expression Omnibus (GEO) as GSE149040 and GSE139904. Bulk RNA-seq data have been deposited at GEO as GSE332728 and are publicly available as of the date of publication. All original code has been deposited at https://github.com/hlbushnell/bushnell-retina and is publicly available as of the date of publication.

## ACKNOWLEDGMENTS

The authors would like to thank Ryan Delgado, Rachel Savage, Ryoji Amamoto, Emma West, and ChangHee Lee for their expertise and fruitful discussions of this work. We also thank the Microscopy Resources on the North Quad (MicRoN) at HMS for their training and microscope maintenance.

## FUNDING

This study was funded by the Howard Hughes Medical Institute and the National Institutes of Health Visual Neuroscience training grant.

## AUTHOR CONTRIBUTIONS

All conceptualization, investigation, and writing by H.L.B. and C.C.

## COMPETING INTERESTS

The authors declare no competing interests.

## MATERIALS AND METHODS

### Mice

All animal procedures were performed in accordance with institutional guidelines and approved by the Institutional Animal Care and Use Committee (IACUC) at Harvard Medical School. Mice were housed under a 12hr light/dark cycle with ad libitum access to food and water. Both male and female embryos and pups were used for all experiments. A CD-1 IGS background (Charles River, Strain #022) was used for all wild type mice in this study due to their large litter sizes. The day of vaginal plug detection was designated as embryonic day 0.5 (E0.5) or was determined by Charles River. The day of birth was designated as P0. Genetically modified mice used in this study include Gal-IRES-Cre (from the lab of Dr. Heike Müenzberg-Gruening; (Laque et al., 2015)), Ai140D (The Jackson Laboratory, Strain #030220; (Daigle et al., 2018)), R26R-Confetti (The Jackson Laboratory, Strain #017492; (Snippert et al., 2010)), and Mcm2-CreERT2 (The Jackson Laboratory, strain #017611; (Pruitt et al., 2007)). Genetic crosses were performed to obtain mice with the genotypes: Gal-Cre/+; Ai140/+, Gal-Cre/+; Confetti/+, Mcm2-CreERT2/+; Ai140/+.

### scRNA-seq dataset processing and integration

Two published scRNA-seq datasets from embryonic mouse retina were obtained from NCBI GEO and processed in R (v4.5.0) using Seurat v5.3.1. The first dataset (Wu et al., 2021; GSE149040) contains E14.5 mouse retinal cells; the heterozygous condition (GSM4488285), representing wild-type retinal development, was used for analysis. The second dataset (Balasubramanian et al., 2021; GSE139904) contains E13.5 mouse retinal cells from a conditional Fgfr1/2 double-knockout study in which knockout cells are labeled with a TdTomato reporter. Only TdTomato-negative (wild-type) cells were retained for downstream analysis. Quality control metrics were calculated for each dataset independently, and filtering thresholds for each were determined by ranked gene count distributions. For the Wu et al. dataset, cells were retained with 692 < nFeature_RNA < 3,981 and percent.mt < 4%. Doublets were detected using scDblFinder and 247 putative doublets were removed. After QC filtering, 4,026 cells were retained from an initial 4,639. For the Balasubramanian et al. dataset, cells were retained with 1,778 < nFeature_RNA < 5,623 and percent.mt < 12%. TdTomato-positive cells were removed, and scDblFinder was used to detect and remove 276 doublets from the remaining cells. After QC filtering and TdTomato exclusion, 4,285 cells were retained from an initial 10,671.

The two datasets were merged and integrated using Seurat v5.3.1. Data were log-normalized, and the top 2,000 highly variable features were identified prior to scaling and PCA. Batch effects between datasets were corrected using Canonical Correlation Analysis (CCA) via Seurat’s IntegrateLayers function, yielding a corrected low-dimensional embedding. UMAP dimensionality reduction, nearest-neighbor graph construction, and graph-based clustering (resolution = 1.5) were performed using the first 30 CCA dimensions. A random seed of 42 was used throughout. Non-retinal contaminating clusters (RPE, fibroblast, immune cells) and low-quality cells were identified and removed based on marker gene expression and nFeature_RNA. Following contaminant removal, 7,016 retinal cells were retained. Normalization, variable feature selection, and dimensionality reduction were repeated on the filtered dataset, and final clustering was performed at resolution 0.2 yielding 7 broad clusters. RGC and RPC subclusters were merged to result in 5 final cell class clusters.

### Neurogenic progenitor cell subclustering and differential expression analysis

NPCs were isolated by subsetting the neurogenic cluster from the integrated retinal dataset, identified by co-expression of *Top2a* and *Atoh7*. The neurogenic subset was re-normalized using log-normalization, and the top 2,000 highly variable features were re-identified prior to scaling. UMAP dimensionality reduction and nearest-neighbor graph construction were performed using the first 30 CCA dimensions from the previously computed integration. Initial graph-based clustering at resolution 0.7 yielded 8 clusters. One cluster was identified as post-mitotic cells lacking Top2a expression and was excluded. Following removal, 772 cells were retained. The filtered neurogenic subset underwent a second round of normalization, variable feature selection, scaling, UMAP, and graph-based clustering using identical parameters. Clustering at resolution 0.7 yielded 6 subclusters, which were manually consolidated into three neurogenic populations based on inspection of known marker gene expression: NG1 (subclusters 1 and 4; S-phase marker *Ung*^+^), NG2 (subclusters 0 and 2; *Olig2^+^*), and NG3 (subclusters 3 and 5; *Gal*^+^). Differentially expressed genes between NG2 and NG3 were identified using Seurat’s FindMarkers function (Wilcoxon rank-sum test), with a minimum expression fraction of 0.25 and a log₂ fold-change threshold of 0.25.

### Tissue preparation and sectioning

Embryos were harvested at E14.5 in PBS and whole heads were fixed for 2 hours in 4% paraformaldehyde. After fixation, heads were washed 3 x 5 minutes in PBS, then transferred to 30% sucrose in PBS overnight. Next, they were transferred to a 1:1 solution of [30% sucrose in PBS]:[OCT] for 2 hours. Heads were then frozen in cryomolds and stored at -80°C. Similarly, P21 mice were euthanized, and retinas were dissected and then fixed for 30 minutes in 4% paraformaldehyde. After fixation, retinas were washed 3 x 5 minutes in PBS, then transferred to 7% sucrose for 10 minutes, and transferred to a 1:1 [30% sucrose in PBS]:[OCT] for 1 hour.

Retinas were then frozen in cryomolds and stored at -80°C. 25µm cryosections were thaw-mounted onto Superfrost Plus microscope slides (ThermoFisher, cat# 22-037-246) and dried at room temperature for 30 minutes before storage at -80°C or immediate use. Before use with either HCR RNA-FISH or immunostaining, slides were washed 3 × 5 minutes in PBS with 0.1% Tween-20 (PBSTw) to permeabilize tissue and remove OCT.

### HCR RNA-FISH

Sections were prepared as described above, then HCR was performed following the protocol provided by Molecular Instruments using purchased Probe Hybridization Buffer (PHB), Probe Wash Buffers (PWB), and Amplification Buffer. All probes used in this study were custom designed and purchased from Molecular Instruments (See Supplemental Table 1). All steps were performed in a dark, humidified chamber with 200 µL volumes and HybriSlip Hybridization Covers. Briefly, slides were pre-incubated with PHB at 37°C for 10 minutes, then probe solution (0.4 µL of each probe set per 100 µL PHB) was applied and incubated overnight at 37°C. The following day, slides were washed sequentially for 15 minutes each at 37°C in 100%, 75%, 50%, and 25% PWB in 5X SSCT, then 5X SSCT alone, followed by a 5-minute wash in 5X SSCT at room temperature. Amplifier hairpins were heated at 95°C for 90 seconds, then snap-cooled in the dark at room temperature for 30 minutes. Simultaneously, slides were equilibrated in Amplification Buffer for 30 minutes at room temperature. Hairpin mix (2 µL of each hairpin per 100 µL Amplification Buffer) was applied and incubated overnight in the dark, followed by washes of 2 × 30 minutes then 1 × 5 minutes in 5X SSCT. When applicable, slides were stained with 1 µg/mL DAPI in PBSTw for 1 hour at room temperature. When HCR was combined with immunostaining, HCR was performed first followed by immunostaining as described, except for N1ICD staining which requires antigen retrieval and was therefore performed prior to HCR.

### Antigen retrieval and immunostaining

For antigen retrieval, which was only performed prior to anti-N1ICD immunostaining, slides were washed 3 × 5 minutes in TN Buffer (0.1M Tris-HCl pH 7.5, 0.15M NaCl), placed in a slide box filled with boiling pH 6.0 citrate buffer, and incubated for 7 minutes in an electric vegetable steamer. The slide box was submerged in ice for 15 minutes, after which slides were washed 2 × 5 minutes in TN Buffer followed by 2 × 5 minutes in PBSTw before proceeding with immunostaining as described below.

Sections were prepared as described above. Antibodies used in this study were anti-Cleaved Notch1 (Val1744) (Cell Signaling Technology, Cat#4147S, 1:100), anti-Melanopsin (N-terminus) (from Dr. Michael Do, 1:1000), anti-Vsx2 (Chx10) (ThermoFisher, Cat#PA1-12565). Slides were incubated with primary antibody in PBSTw with 5% ChemiBLOCKER (MilliporeSigma Cat#2170) at 4°C overnight, except anti-N1ICD and anti-Melanopsin, which were incubated for 48 and 72 hours, respectively. Slides were washed 5 x 10 minutes with PBSTw, then incubated overnight at 4°C with secondary antibody (1:300) in PBSTw with 3% ChemiBLOCKER and 1µg/mL DAPI. Slides were then washed 3 x 5 minutes in PBSTw prior to mounting.

### EdU labeling

A 10mM EdU working solution was prepared by dissolving 5mg EdU from the Click-iT Plus kit (ThermoFisher Cat#C10637) in 2mL of PBS, with brief warming until EdU is fully dissolved. 350uL of 10mM EdU solution was administered by intraperitoneal injection into E12.5 pregnant dams with an insulin syringe 20 minutes before euthanasia and tissue collection. Incorporated EdU was detected using the provided Click-iT Plus protocol with shortened incubation time (15 minutes) to minimize sample degradation.

### Expression history labeling and cell type quantification

To label cells with *Galanin* expression history, Gal-Cre mice were crossed with Ai140 reporter mice to generate Gal-Cre; Ai140/+ offspring. Mice were genotyped as pups to confirm the presence of both alleles. At P21, retinas were dissected and processed as described above. One retina from each of 5 mice was used for analysis. HCR RNA-FISH was performed using the marker gene combinations described in Supplemental Figure 4, and images were acquired using a 20x objective. Per retina, 5 fields of view were captured from the central third of central retinal sections.

Cell nuclei were segmented using Cellpose-SAM (Pachitariu et al. 2025), run on the DAPI channel with a custom model trained on 3 manually corrected segmentations. Segmentation masks were imported into Napari (www.napari.org; Sofroniew et al. 2022) where cells were manually assigned to cell classes based on HCR RNA-FISH fluorescence and EGFP positivity. Annotations were exported as .csv files and combined for downstream analysis. Statistical comparisons between total and EGFP^+^ cell class proportions were performed using paired two-tailed t-tests in GraphPad Prism (v11.0.1).

To identify pairs of cells produced by division of a *Galanin*-expressing progenitor cell, Gal-Cre mice were crossed with R26R-Confetti reporter mice to generate Gal-Cre; Confetti/+ offspring, genotyped as above. At P21, retinas were dissected and processed as described above, with each retina sectioned entirely onto a single slide. One retina from each of 5 mice was used for analysis. HCR RNA-FISH was performed as described above using probes targeting *Slc32a1*. Retinal sections were screened for same-color, radially aligned cell pairs. Only same-color cells separated by ≤1 soma diameter (∼10 µm) were scored as putative clonal pairs. The first 30 clonal pairs encountered per retina were recorded (150 total; 2 excluded for technical reasons, final n = 148). Cell classes were assigned based on cell body position, morphology, and *Slc32a1* positivity. For identification of ipRGCs, anti-Melanopsin immunostaining was performed on sections from 2 additional Gal-Cre; Confetti/+ retinas.

### Embryonic clonal labeling

To sparsely label individual progenitor cells, Mcm2-CreERT2/+ mice were crossed with Ai140 reporter mice to generate Mcm2-CreERT2/+; Ai140/Ai140 offspring. Timed matings were performed between male Mcm2-CreERT2/+; Ai140/Ai140 mice and female CD-1 mice, which were used for their large litter sizes. E12.5 dams were administered 400 µL of 4 mg/mL tamoxifen (Sigma-Aldrich, Cat#T5648) in corn oil by oral gavage. Dams were euthanized 26-40 hours later to allow labeled progenitor cells to divide at least once following Cre-mediated reporter activation. Embryos were harvested and Mcm2-CreERT2/+; Ai140/+ embryos were identified by EGFP fluorescence throughout the body under a dissecting microscope. Embryonic tissue was collected and processed as described above. Retinal sections were screened by HCR RNA-FISH for *Gal* and *Olig2* to identify two-cell clones containing at least one *Gal*+ or *Olig2*+ cell. Given the sparseness of labeled clones and the rarity of NPC-containing clones, retinas from multiple dams were used (n = 10 retinas, 90 clones total). To identify the clonal siblings of NPCs, a subset of sections was stained by HCR RNA-FISH for *Gal* or *Olig2* combined with anti-Vsx2 immunostaining.

### Single cell quantification of HCR RNA-FISH and immunofluorescence

Embryonic retinal sections were prepared and imaged as described above. Nuclear fluorescence intensities were quantified using a custom Python script. Briefly, nuclear segmentation masks generated by Cellpose-SAM (as described above) were used to define nuclear regions of interest. Mean fluorescence intensity within each nucleus was extracted for each channel using the regionprops_table function from scikit-image (van der Walt et al., 2014). Per-nucleus intensity values were exported as .csv files for statistical analysis in GraphPad Prism (v11.0.1).

### Embryonic retinal electroporation and ex vivo culture

Following euthanasia of pregnant dams, uterine horns were dissected and placed in room temperature PBS. Embryos were removed, decapitated, and whole eyes were dissected with forceps and transferred to a dissection dish containing PBS. Plasmid DNA (0.5 µg/µL, per plasmid, in 1X PBS with 0.01% Fast Green) was injected subretinally using a pulled glass needle. All retinas were electroporated with a bpCAG-EGFP expression plasmid to label electroporated cells. For Notch signaling activation, retinas were co-electroporated with a bpCAG-NICD expression plasmid. Injected eyes were electroporated using a square-wave electroporator with tweezer-type electrodes delivering five 50 ms pulses of 40 V at 1-second intervals. Electroporated retinas were then dissected from whole eye globes, retaining the lens for structural support.

Retinas were cultured ex vivo for 16 hours in 1 mL of culture medium consisting of a 1:1 mixture of DMEM (ThermoFisher, Cat#11965118) and Ham’s F-12 (ThermoFisher, Cat#11765054), supplemented with 10% FBS and Pen/Strep. Retinas were cultured in 1.5 mL Eppendorf tubes with gentle rocking at 37°C at a density of no more than 8 retinas per tube. For Notch signaling inhibition, LY411575 (Sigma-Aldrich, Cat#SML0506) was added to culture medium at a final concentration of 1 µM. Following culture, retinas were either fixed and processed as described above, or dissociated and subjected to FACS isolation of EGFP^+^ cells as described below.

### Dissociation and fluorescence-activated cell sorting (FACS)

Embryonic retinas were dissociated using the Worthington Papain Dissociation System (Worthington, Cat# LK003150). Papain solution and Ovomucoid solution were prepared according to the manufacturer’s instructions, with the addition of 15 µL/mL 1M HEPES buffer and 1 µL/mL RNase inhibitor (ThermoFisher, Cat# Am2694) to the Papain solution. Papain solution was pre-warmed at 37°C for at least 10 minutes prior to use. Explant culture medium was removed and retinas were washed with 1 mL PBS, then transferred to a 5 mL polypropylene tube containing 1 mL pre-warmed Papain solution and incubated at 37°C for 30 minutes with gentle agitation every ∼5 minutes. Retinas were mechanically dissociated by trituration with a P1000 pipette, after which 1 mL Ovomucoid solution was added and gently mixed. Cells were pelleted by centrifugation at 360 x g for 5 minutes, the supernatant was removed, and the pellet was resuspended in 300 µL of running buffer (PBS with 1% BSA, 1 µL/mL RNase inhibitor, and 2 µL/mL 0.5M EDTA). Cell suspensions were filtered through a 35 µm cell strainer cap into a polypropylene FACS tube prior to sorting.

EGFP+ cells were sorted on a BD FACSAria III cell sorter using an 85 µm nozzle. Cells were sorted directly into 500 µL of collection buffer (PBS with 1 µL/mL RNase inhibitor and 2 µL/mL DNase (included in Dissociation System)) in a Lo-Bind 1.5 mL Eppendorf tube. Sorted cells were then processed for RNA extraction or bulk RNA sequencing as described below.

### Bulk RNA sequencing

Following FACS, sorted cells were pelleted by centrifugation at 600 x g for 5 minutes. The supernatant was removed to a remaining volume of 25 µL, and 25 µL of 2x DNA/RNA Shield (Zymo Research, cat# R1200) was added and gently mixed. Samples containing approximately 60,000-150,000 EGFP^+^ cells were submitted to Plasmidsaurus for total RNA extraction and library preparation. Libraries were prepared using a 3’ end counting approach: mRNA was converted to cDNA via reverse transcription using a poly(dT)VN primer, followed by second-strand synthesis, tagmentation, library indexing, and amplification. Unique molecular identifiers (UMIs) and unique dual indices (UDIs) were incorporated to enable PCR deduplication and prevent index hopping, respectively. Libraries were sequenced on an Illumina platform with single-end 90 bp reads, targeting approximately 10 million deduplicated reads per sample.

Raw reads were processed by Plasmidsaurus using their standard pipeline: quality filtering and adapter trimming with FastP (v0.24.0; minimum Phred quality score of 15, minimum read length 50 bp), alignment to the mouse reference genome (GRCm39) using STAR (v2.7.11), UMI-based deduplication using UMICollapse (v1.1.0), and gene expression quantification using featureCounts (subread v2.1.1) with Ensembl release 114 annotation. Gene count tables were used for downstream differential expression analysis as described in Quantification and Statistical Analysis. Raw sequencing data have been deposited in the NCBI Gene Expression Omnibus (GEO).

### RT-Droplet Digital PCR (RT-ddPCR)

Total RNA was extracted from sorted EGFP+ cells preserved in TRIzol LS Reagent (ThermoFisher, Cat# 10296028) using the Zymo Direct-zol RNA Miniprep kit (Zymo Research, Cat# R2052) according to the manufacturer’s instructions. RNA concentration was measured by spectrophotometry (NanoDrop), and cDNA was synthesized from the full extracted volume using LunaScript RT SuperMix (NEB, Cat# M3010L). To normalize cDNA input across samples, cDNA was diluted to a common concentration based on the least concentrated sample, and a volume corresponding to 3 ng of input RNA was added to each ddPCR reaction.

ddPCR was performed using the Bio-Rad QX200 AutoDG Droplet Digital PCR System with ddPCR EvaGreen Supermix (Bio-Rad, cat# 1864034). Each 20 µL reaction contained cDNA, 1x EvaGreen Supermix, and 250 nM each of forward and reverse primers targeting *Dll1*, *Dll4*, or *Hprt* (primer sequences provided in Key Resources Table). Droplets were generated automatically using the AutoDG system, thermocycled according to the manufacturer’s recommended EvaGreen protocol, and read on the QX200 reader. Data were analyzed using QuantaSoft software.

Expression of *Dll1* and *Dll4* was normalized to *Hprt* as a housekeeping gene and reported as log2 fold change relative to the mean expression in the GFP-only control condition. Statistical analysis was performed as described in Quantification and Statistical Analysis.

### Image acquisition

All slides were mounted with Fluoromount-G prior to imaging. Samples were imaged with a Yokogawa CSU-W1 single disk (50mm pinhole size) spinning disk confocal unit attached to a fully motorized Nikon Ti2 inverted microscope equipped with a Nikon linear-encoded motorized stage with a Mad City Labs 500µm range Nano-Drive Z piezo insert, an Andor Zyla 4.2 plus (6.5mm photodiode size) sCMOS camera using a Nikon ApolS LWD 40X/1.1 DICN2 water immersion objective lens with Zeiss Immersol W 2010. The final digital resolution of the image was 0.16µm/pixel. Fluorescence from (405 nm, 488 nm, 550 nm, and 640 nm) was collected by illuminating the sample with directly modulated solid-state lasers 405 nm diode 100mW (at the fiber tip) laser line, 488 nm diode 100mW laser line, 561nm DPSS 100mW laser line and 640 nm diode 70mW laser line in a Toptica iChrome MLE laser combiner, respectively. A hard-coated Semrock Di01-T405/488/568/647 multi-bandpass dichroic mirror was used for all channels. Signal from each channel was acquired sequentially with hard-coated Chroma ET455/50, Chroma ET525/36nm, Chroma ET605/52nm emission filters and Chroma ET705/72nm in a filter wheel placed within the scan unit, for blue, green, red and far-red channels, respectively. All images were captured using the 16-bit dual gain high dynamic range camera mode and no binning. Nikon Elements AR 5.02 acquisition software was used to acquire the data. Z-stacks were acquired using Piezo Z-device, with the shutter closed during axial movement. Images were acquired by collecting the entire Z-stack in each color. Data were saved as ND2 files.

### Quantification and Statistical Analysis

Data analysis and visualization were performed using R software (v.4.5.0) and GraphPad Prism (version 11.0.0). Statistical analyses, including significance values, error bars, number of replicates and mice per group, and scale bars, are indicated within the figures and their legends. Specifically, R packages such as Seurat (v.5.3.1) and DESeq2 were used for statistical analysis. Statistical significance was determined using paired Student’s *t*-test in analysis comparing two conditions, two-way ANOVA for grouped analysis, and Kruskal-Wallis test with Dunn’s post-hoc for multiple pairwise comparisons.

### AI Usage

During the preparation of this work, the author(s) used GPT-5.2 and Claude 4.6 to write and de-bug code as well as edit the manuscript for spelling, grammar, and clarity. After using this tool or service, the authors reviewed and edited the content as needed and take full responsibility for the content of the publication.

---

**Supplemental Fig. 1.**
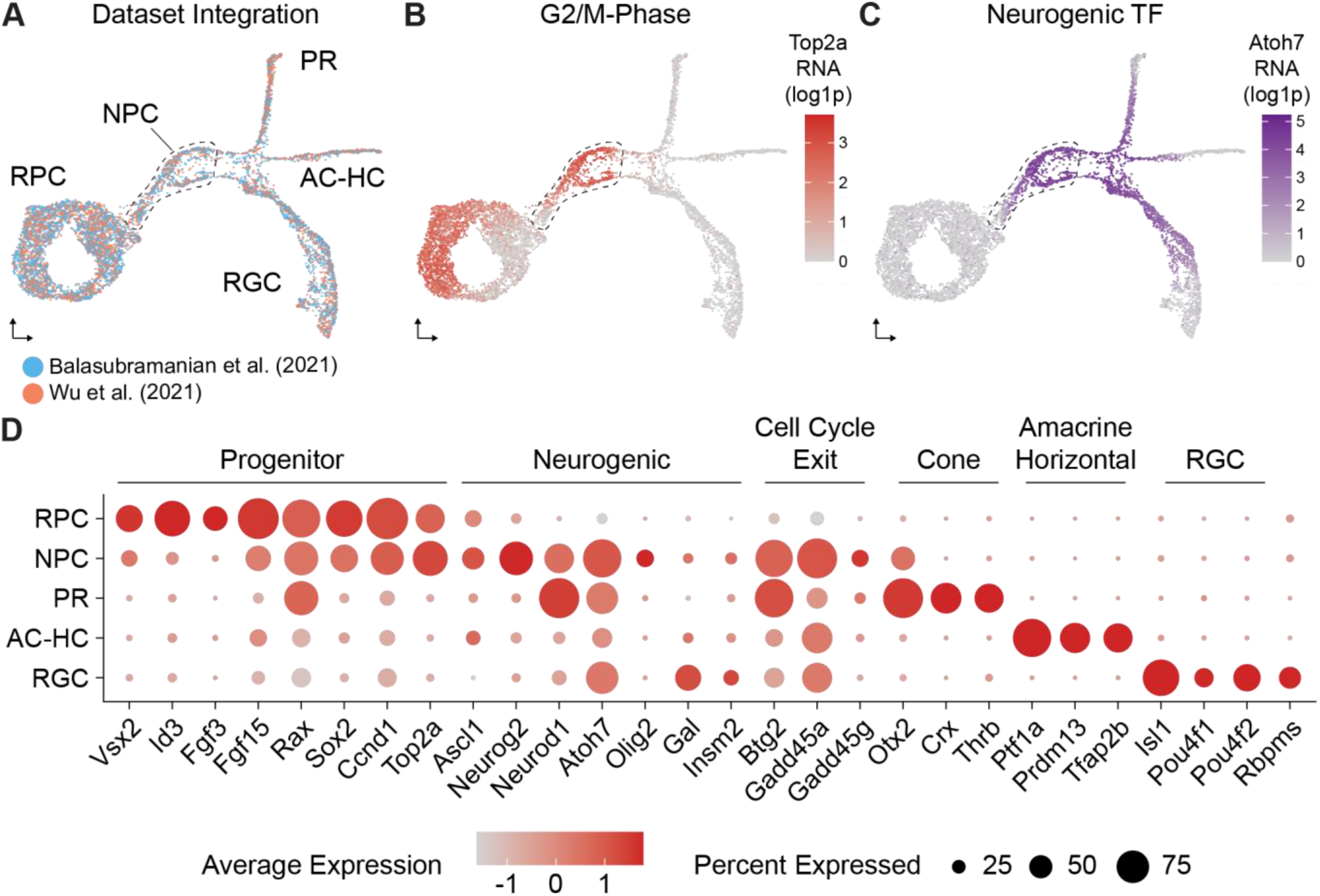
Gene expression from integrated scRNA-seq data. A) UMAP of integrated E13.5-E14.5 mouse retina scRNA-seq (n = 7,258 cells). Cells colored by dataset of origin. B) UMAP as in A, with cells colored by expression of G2/M marker Top2a. Dotted line indicates presumptive NPCs. C) UMAP as in A, with cells colored by expression of NPC marker Atoh7. D) Dot plot showing aggregate gene expression of marker genes for each broad cell class identified in Figure 1B, including progenitor markers, neurogenic markers, and cell cycle exit factors. Dot size indicates the percentage of cells expressing each gene; color intensity reflects mean expression scaled across clusters (z-score).

**Supplemental Fig. 2.**
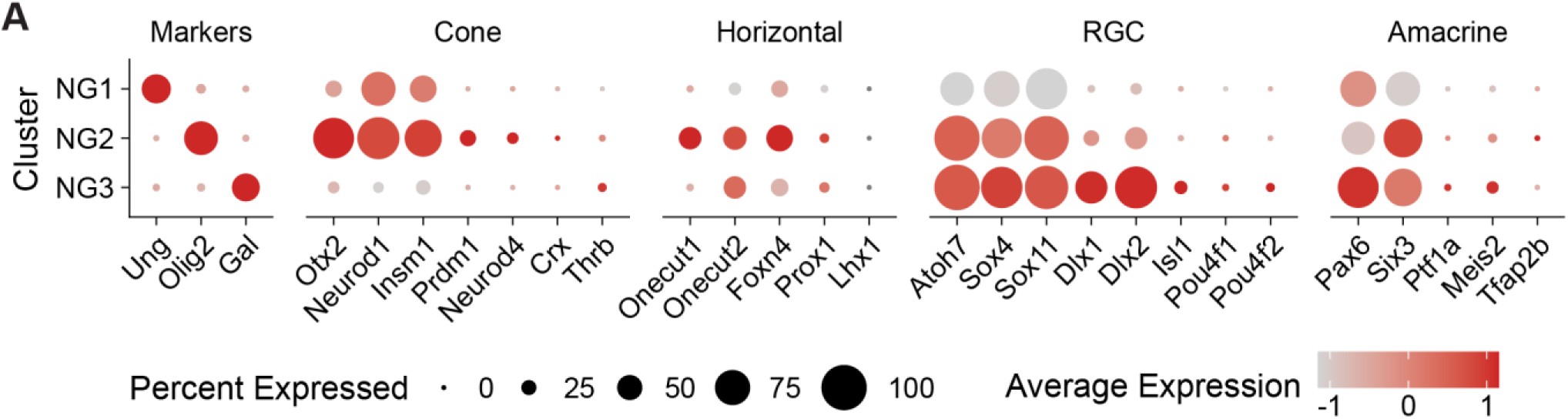
Gene expression from neurogenic subclusters. A) Dot plot showing aggregate gene expression in cells from each neurogenic subcluster identified in Figure 1C, highlighting genes associated with distinct post-mitotic retinal cell classes. As in Supp. Fig. 1D, color reflects scaled expression.

**Supplemental Fig. 3.**
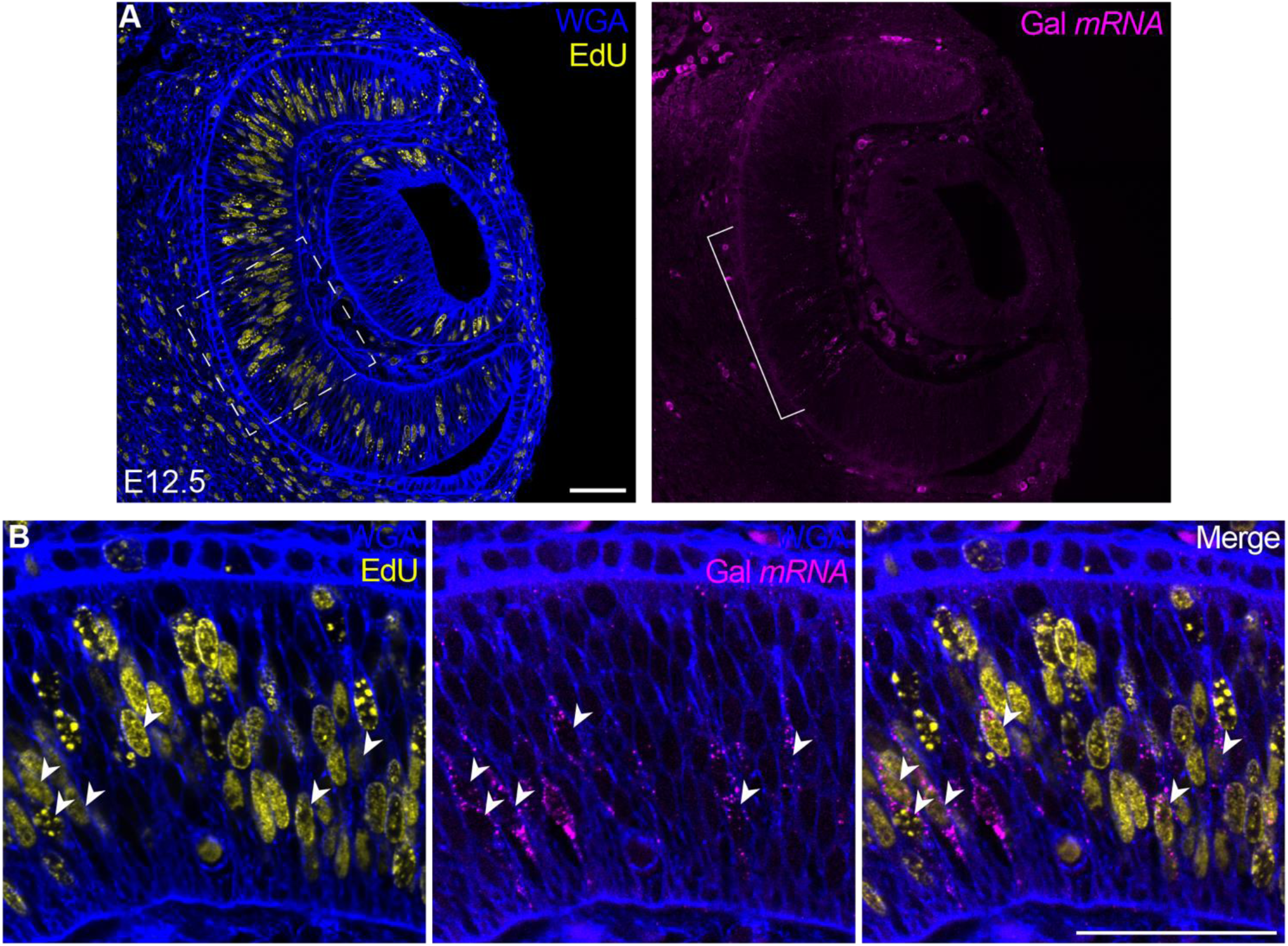
*Gal*^+^ NPCs restricted to central retina at E12.5. (A) Representative image of E12.5 retinal cross section with EdU labeling (yellow) and HCR RNA-FISH for *Gal* (magenta). WGA (blue) marks cell boundaries. Scale bar, 50µm. Bracket indicates region containing *Gal*^+^ cells. Dotted line indicates boundary of the region in (B). (B) Higher magnification image of central region from (A). Arrows indicate EdU^+^/Gal^+^ double-positive cells. Scale bar, 50µm.

**Supplemental Fig. 4.**
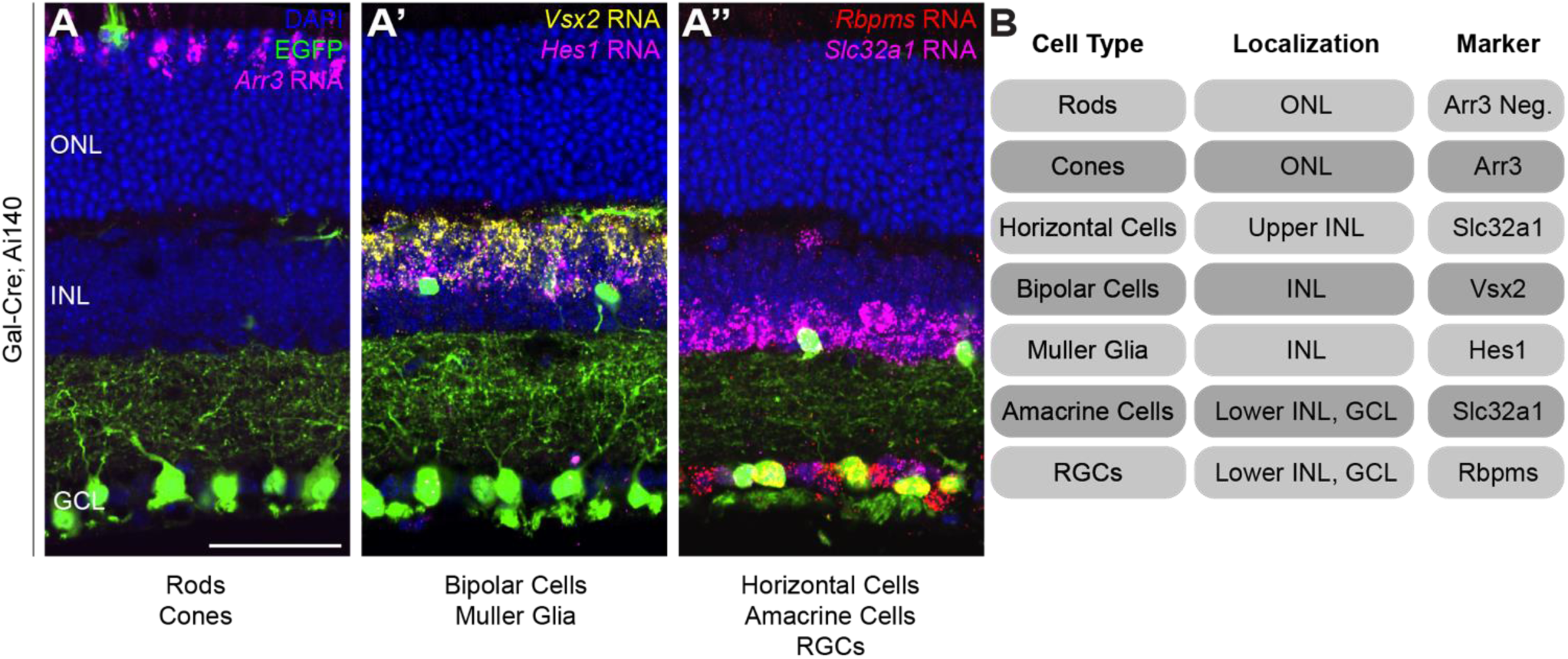
In situ cell type identification. (A-A’’) Example images from P21 Gal-Cre; Ai140 mouse retinas with EGFP (green) and stained for DAPI (blue) with HCR RNA-FISH for cell type markers: (A) Arr3 (magenta), (A’) Vsx2 (yellow) and Hes1 (magenta), (A’’) Rbpms (red) and Slc32a1 (magenta). Scale bar, 25µm (applies to all images). (B) Table of cell body localizations and marker genes used to identify each cell type.

**Supplemental Fig. 5.**
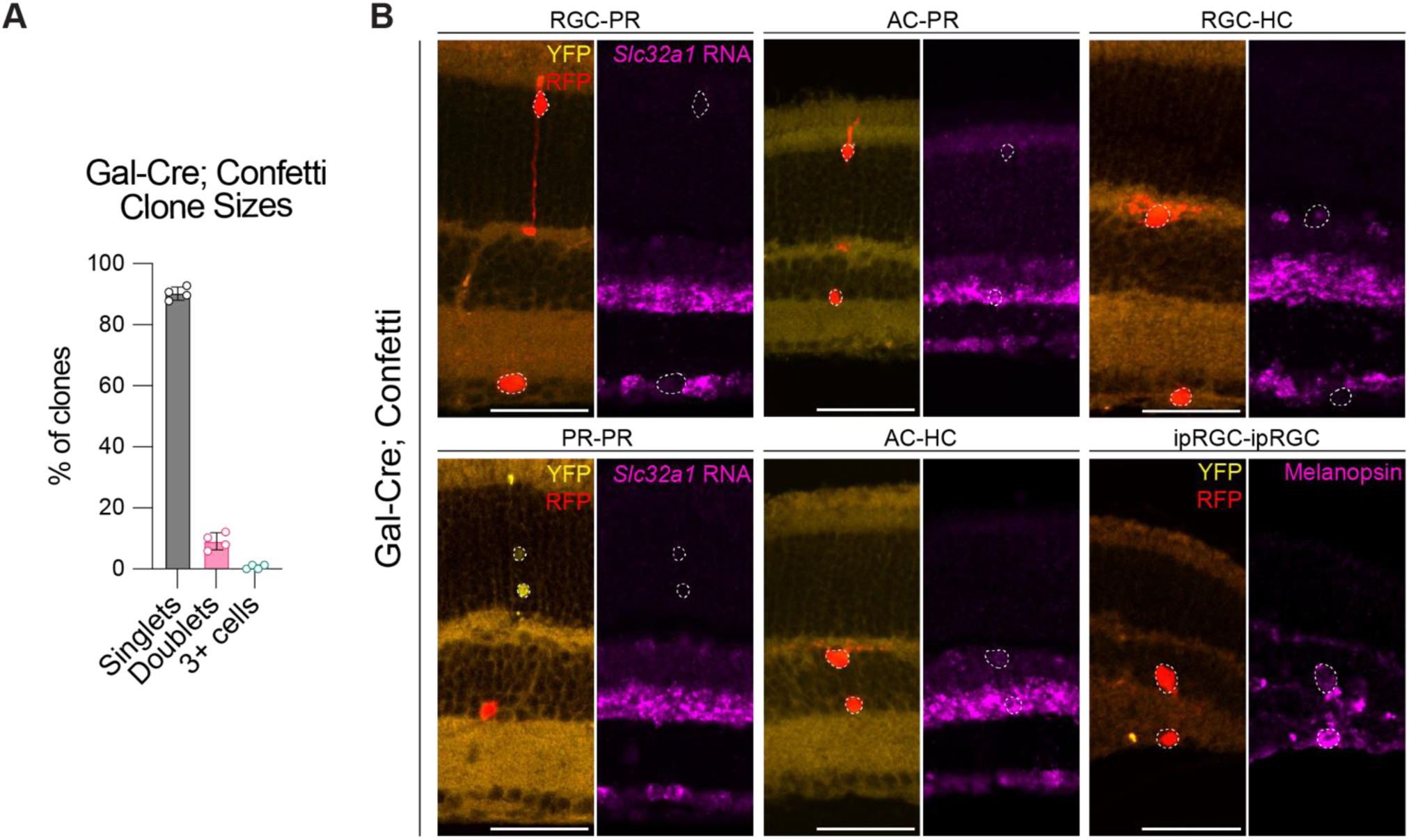
Gal-Cre; Confetti clones. (A) Bar graph showing the distribution of Gal-Cre; Confetti recombination events observed as singlets, doublets, and groups of 3+ cells (n = 4 retinas, one full 30µm section per retina, average ∼18 recombination events per section). (B) Examples of two-cell clone compositions observed at lower frequency and not shown in Figure 2, from p21 Gal-Cre; Confetti mice. For each panel: (Left) YFP (yellow) and RFP (red) from Confetti reporter recombination. (Right) HCR RNA-FISH for amacrine/horizontal cell marker *Slc32a1* (magenta). Scale bars, 50 µm.

**Supplemental Fig. 6.**
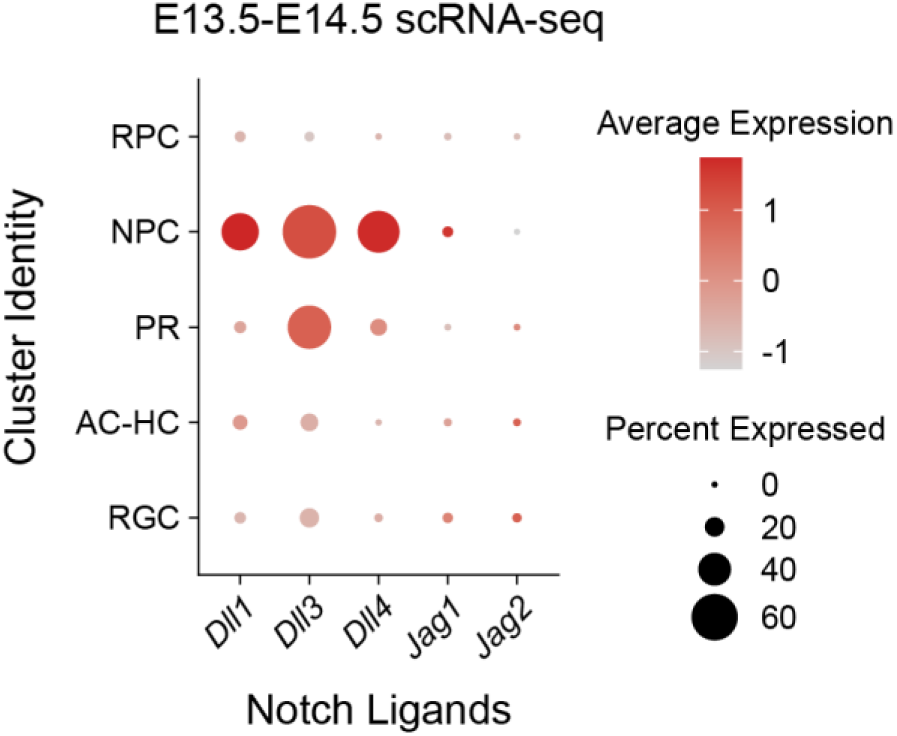
Notch ligand expression in E13.5-E14.5 retina. A) Dot plot showing aggregate expression of Notch ligand genes in cells from each retinal cluster identified in Figure 1B. As in Supp. Fig. 1, color reflects scaled expression.

**Supplemental Table 1.** HCR RNA-FISH probes. Table of custom HCR RNA-FISH probes used in this study, listing the target, source, and identifier of each probe set.

| HCR RNA-FISH Probe Set Target | Source | Identifier |
| --- | --- | --- |
| Mouse *Gal* (RefSeq NM_010253) | Molecular Instruments | Lot #PRC079; HCR v3.0; Amp. B2 |
| Mouse *Olig2* (RefSeq NM_016967) | Molecular Instruments | Lot #RTE112; HCR v3.0; Amp. B3 |
| Mouse *Slc32a1* (RefSeq NM_009508.3) | Molecular Instruments | Lot #RTC591; HCR v3.0; Amp. B2 |
| Mouse *Rbpms* (RefSeq NM_001042674.2) | Molecular Instruments | Lot #RTJ837; HCR v3.0; Amp. B3 |
| Mouse *Dll1* (RefSeq NM_007865.3) | Molecular Instruments | Lot #PRN917; HCR v3.0; Amp. B4 |
| Mouse *Dll4* (RefSeq NM_019454.4) | Molecular Instruments | Lot #PRN918; HCR v3.0; Amp. B5 |
| Mouse *Arr3* (RefSeq NM_133205.3) | Molecular Instruments | HCR v3.0; Amp. B1 |
| Mouse *Vsx2* (RefSeq NM_001301427.1) | Molecular Instruments | Lot #RTU599; HCR v3.0; Amp. B4 |
| Mouse *Hes1* (RefSeq BC018375.1) | Molecular Instruments | Lot #PRH379; HCR v3.0; Amp. B1 |

## Notes

### Competing Interest Statement

The authors have declared no competing interest.

## REFERENCES

1. Austin, C. P., Feldman, D. E., Ida, J. A., Jr. and Cepko, C. L. (1995). Vertebrate retinal ganglion cells are selected from competent progenitors by the action of Notch. Development 121, 3637– 3650.

2. Balasubramanian, R., Min, X., Quinn, P. M. J., Giudice, Q. L., Tao, C., Polanco, K., Makrides, N., Peregrin, J., Bouaziz, M., Mao, Y., et al. (2021). Phase transition specified by a binary code patterns the vertebrate eye cup. Science Advances 7,.

3. Berson, D. M., Dunn, F. A. and Takao, M. (2002). Phototransduction by retinal ganglion cells that set the circadian clock. Science 295, 1070–3.

4. Bosze, B., Moon, M. S., Kageyama, R. and Brown, N. L. (2020). Simultaneous Requirements for Hes1 in Retinal Neurogenesis and Optic Cup-Stalk Boundary Maintenance. J Neurosci 40, 1501–1513.

5. Bosze, B., Suarez-Navarro, J., Cajias, I., Brzezinski Iv, J. A. and Brown, N. L. (2023). Notch pathway mutants do not equivalently perturb mouse embryonic retinal development. PLOS Genetics 19, e1010928.

6. Brodie-Kommit, J., Clark, B. S., Shi, Q., Shiau, F., Kim, D. W., Langel, J., Sheely, C., Ruzycki, P. A., Fries, M., Javed, A., et al. (2021). Atoh7-independent specification of retinal ganglion cell identity. Science Advances 7, eabe4983.

7. Brody, T. and Odenwald, W. F. (2000). Programmed Transformations in Neuroblast Gene Expression during *Drosophila* CNS Lineage Development. Developmental Biology 226, 34–44.

8. Brzezinski, J. A., Kim, E. J., Johnson, J. E. and Reh, T. A. (2011). Ascl1 expression defines a subpopulation of lineage-restricted progenitors in the mammalian retina. Development 138, 3519–3531.

9. Brzezinski, J. A., Prasov, L. and Glaser, T. (2012). Math5 defines the ganglion cell competence state in a subpopulation of retinal progenitor cells exiting the cell cycle. Developmental Biology 365, 395–413.

10. Cepko, C. (2014). Intrinsically different retinal progenitor cells produce specific types of progeny. Nature Reviews Neuroscience 15, 615–627.

11. Chan, M. M., Smith, Z. D., Grosswendt, S., Kretzmer, H., Norman, T. M., Adamson, B., Jost, M., Quinn, J. J., Yang, D., Jones, M. G., et al. (2019). Molecular recording of mammalian embryogenesis. Nature 570, 77–82.

12. Chen, X. and Emerson, M. M. (2021). Notch signaling represses cone photoreceptor formation through the regulation of retinal progenitor cell states. Scientific Reports 11,.

13. Choi, J., Li, J., Ferdous, S., Liang, Q., Moffitt, J. R. and Chen, R. (2023). Spatial organization of the mouse retina at single cell resolution by MERFISH. Nat Commun 14, 4929.

14. Clark, B. S., Stein-O’Brien, G. L., Shiau, F., Cannon, G. H., Davis-Marcisak, E., Sherman, T., Santiago, C. P., Hoang, T. V., Rajaii, F., James-Esposito, R. E., et al. (2019). Single-Cell RNA-Seq Analysis of Retinal Development Identifies NFI Factors as Regulating Mitotic Exit and Late-Born Cell Specification. Neuron 102, 1111–1126.e5.

15. Daigle, T. L., Madisen, L., Hage, T. A., Valley, M. T., Knoblich, U., Larsen, R. S., Takeno, M. M., Huang, L., Gu, H., Larsen, R., et al. (2018). A Suite of Transgenic Driver and Reporter Mouse Lines with Enhanced Brain-Cell-Type Targeting and Functionality. Cell 174, 465–480.e22.

16. Dräger, U. and Olsen, J. F. (1981). Ganglion cell distribution in the retina of the mouse. Investigative ophthalmology & visual science 20, 285–293.

17. Duda, S., Block, C. T., Pradhan, D. R., Arzhangnia, Y., Klaiber, A., Greschner, M. and Puller, C. (2025). Spatial distribution and functional integration of displaced retinal ganglion cells. Scientific Reports 15,.

18. Elliott, J., Jolicoeur, C., Ramamurthy, V. and Cayouette, M. (2008). Ikaros Confers Early Temporal Competence to Mouse Retinal Progenitor Cells. Neuron 60, 26–39.

19. Emerson, M. M., Surzenko, N., Goetz, J. J., Trimarchi, J. M. and Cepko, C. L. (2013). Otx2 and Onecut1 promote the fates of cone photoreceptors and horizontal cells and repress rod photoreceptors. Developmental Cell 26 1, 59–72.

20. Ge, Y., Chen, X., Nan, N., Bard, J., Wu, F., Yergeau, D., Liu, T., Wang, J. and Mu, X. (2023). Key transcription factors influence the epigenetic landscape to regulate retinal cell differentiation. Nucleic Acids Res 51, 2151–2176.

21. Goetz, J., Jessen, Z. F., Jacobi, A., Mani, A., Cooler, S., Greer, D., Kadri, S., Segal, J., Shekhar, K., Sanes, J. R., et al. (2022). Unified classification of mouse retinal ganglion cells using function, morphology, and gene expression. Cell Reports 40, 111040.

22. Gozlan, O. and Sprinzak, D. (2023). Notch signaling in development and homeostasis. Development 150, dev201138.

23. Guido, M. E., Marchese, N. A., Rios, M. N., Morera, L. P., Diaz, N. M., Garbarino-Pico, E. and Contin, M. A. (2022). Non-visual Opsins and Novel Photo-Detectors in the Vertebrate Inner Retina Mediate Light Responses Within the Blue Spectrum Region. Cell Mol Neurobiol 42, 59– 83.

24. Hafler, B. P., Surzenko, N., Beier, K. T., Punzo, C., Trimarchi, J. M., Kong, J. H. and Cepko, C. L. (2012). Transcription factor *Olig2* defines subpopulations of retinal progenitor cells biased toward specific cell fates. Proceedings of the National Academy of Sciences 109, 7882–7887.

25. Hattar, S., Lucas, R. J., Mrosovsky, N., Thompson, S., Douglas, R. H., Hankins, M. W., Lem, J., Biel, M., Hofmann, F., Foster, R. G., et al. (2003). Melanopsin and rod–cone photoreceptive systems account for all major accessory visual functions in mice. Nature 424, 75–81.

26. He, J., Zhang, G., Almeida, A. D., Cayouette, M., Simons, B. D. and Harris, W. Variable Clones Build an Invariant Retina. Neuron 75, 786–798.

27. Heitzler, P. and Simpson, P. (1991). The choice of cell fate in the epidermis of Drosophila. Cell 64, 1083–1092.

28. Henrique, D., Hirsinger, E., Adam, J., Roux, I. L., Pourquié, O., Ish-Horowicz, D. and Lewis, J. (1997). Maintenance of neuroepithelial progenitor cells by Delta–Notch signalling in the embryonic chick retina. Current Biology 7, 661–670.

29. Isshiki, T., Pearson, B., Holbrook, S. and Doe, C. Q. (2001). Drosophila Neuroblasts Sequentially Express Transcription Factors which Specify the Temporal Identity of Their Neuronal Progeny. Cell 106, 511–521.

30. Ivanov, D. (2019). Notch Signaling-Induced Oscillatory Gene Expression May Drive Neurogenesis in the Developing Retina. Front. Mol. Neurosci. 12,.

31. Jadhav, A. P., Cho, S.-H. and Cepko, C. L. (2006a). Notch activity permits retinal cells to progress through multiple progenitor states and acquire a stem cell property. Proceedings of the National Academy of Sciences 103, 18998–19003.

32. Jadhav, A. P., Mason, H. A. and Cepko, C. L. (2006b). Notch 1 inhibits photoreceptor production in the developing mammalian retina. Development 133, 913–923.

33. Jeon, C. J., Strettoi, E. and Masland, R. H. (1998). The major cell populations of the mouse retina. J Neurosci 18, 8936–46.

34. Kageyama, R., Ohtsuka, T., Shimojo, H. and Imayoshi, I. (2008). Dynamic Notch signaling in neural progenitor cells and a revised view of lateral inhibition. Nature Neuroscience 11, 1247–1251.

35. Kaufman, M. L., Park, K. U., Goodson, N. B., Chew, S., Bersie, S., Jones, K. L., Lamba, D. A. and Brzezinski, J. A. (2019). Transcriptional profiling of murine retinas undergoing semi-synchronous cone photoreceptor differentiation. Developmental Biology 453, 155–167.

36. Laque, A., Yu, S., Qualls-Creekmore, E., Gettys, S., Schwartzenburg, C., Bui, K., Rhodes, C., Berthoud, H.-R., Morrison, C. D., Richards, B. K., et al. (2015). Leptin modulates nutrient reward via inhibitory galanin action on orexin neurons. Molecular Metabolism 4, 706–717.

37. Lee, H. Y., Wroblewski, E., Philips, G. T., Stair, C. N., Conley, K., Reedy, M., Mastick, G. S. and Brown, N. L. (2005). Multiple requirements for Hes 1 during early eye formation. Dev Biol 284, 464–78.

38. Lemaire, P. (2009). Unfolding a chordate developmental program, one cell at a time: Invariant cell lineages, short-range inductions and evolutionary plasticity in ascidians. Developmental Biology 332, 48–60.

39. Louvi, A. and Artavanis-Tsakonas, S. (2006). Notch signalling in vertebrate neural development. Nat Rev Neurosci 7, 93–102.

40. Luo, H., Jin, K., Xie, Z., Qiu, F., Li, S., Zou, M., Cai, L., Hozumi, K., Shima, D. T. and Xiang, M. (2012). Forkhead box N4 (Foxn4) activates Dll4-Notch signaling to suppress photoreceptor cell fates of early retinal progenitors. Proceedings of the National Academy of Sciences 109, E553– E562.

41. Manning, C. S., Biga, V., Boyd, J., Kursawe, J., Ymisson, B., Spiller, D. G., Sanderson, C. M., Galla, T., Rattray, M. and Papalopulu, N. (2019). Quantitative single-cell live imaging links HES5 dynamics with cell-state and fate in murine neurogenesis. Nature Communications 10,.

42. Mao, C.-A., Li, H., Zhang, Z., Kiyama, T., Panda, S., Hattar, S., Ribelayga, C. P., Mills, S. L. and Wang, S. W. (2014). T-box Transcription Regulator Tbr2 Is Essential for the Formation and Maintenance of Opn4/Melanopsin-Expressing Intrinsically Photosensitive Retinal Ganglion Cells. J. Neurosci. 34, 13083–13095.

43. Maslov, A. Y., Bailey, K. J., Mielnicki, L. M., Freeland, A. L., Sun, X., Burhans, W. C. and Pruitt, S. C. (2007). Stem/Progenitor Cell-Specific Enhanced Green Fluorescent Protein Expression Driven by the Endogenous Mcm2 Promoter. Stem Cells 25, 132–138.

44. Matsuda, T. and Cepko, C. L. (2004). Electroporation and RNA interference in the rodent retina in vivo and in vitro. Proc Natl Acad Sci U S A 101, 16–22.

45. Mills, E. A. and Goldman, D. (2017). The Regulation of Notch Signaling in Retinal Development and Regeneration. Curr Pathobiol Rep 5, 323–331.

46. Mizeracka, K., DeMaso, C. R. and Cepko, C. L. (2013). Notch1 is required in newly postmitotic cells to inhibit the rod photoreceptor fate. Development 140, 3188–3197.

47. Nelson, B. R., Hartman, B. H., Georgi, S. A., Lan, M. S. and Reh, T. A. (2007). Transient inactivation of Notch signaling synchronizes differentiation of neural progenitor cells. Dev Biol 304, 479–98.

48. Nerli, E., Rocha-Martins, M. and Norden, C. (2020). Asymmetric neurogenic commitment of retinal progenitors involves Notch through the endocytic pathway. eLife 9, e60462.

49. Nerli, E., Kretzschmar, J., Bianucci, T., Rocha-Martins, M., Zechner, C. and Norden, C. (2023). Deterministic and probabilistic fate decisions co-exist in a single retinal lineage. The EMBO Journal 42,.

50. Panda, S., Provencio, I., Tu, D. C., Pires, S. S., Rollag, M. D., Castrucci, A. M., Pletcher, M. T., Sato, T. K., Wiltshire, T., Andahazy, M., et al. (2003). Melanopsin is required for non-image-forming photic responses in blind mice. Science 301, 525–7.

51. Pijuan-Sala, B., Guibentif, C. and Göttgens, B. (2018). Single-cell transcriptional profiling: a window into embryonic cell-type specification. Nat Rev Mol Cell Biol 19, 399–412.

52. Pruitt, S. C., Bailey, K. J. and Freeland, A. (2007). Reduced Mcm2 expression results in severe stem/progenitor cell deficiency and cancer. Stem Cells 25, 3121–32.

53. Reese, B. E., Harvey, A. R. and Tan, S. S. (1995). Radial and tangential dispersion patterns in the mouse retina are cell-class specific. Proceedings of the National Academy of Sciences 92, 2494– 2498.

54. Riesenberg, A. N. and Brown, N. L. (2016). Cell autonomous and nonautonomous Delta-like1 during early mouse retinal neurogenesis. Dev Dyn 245, 631–40.

55. Riesenberg, A. N., Liu, Z., Kopan, R. and Brown, N. L. (2009). Rbpj cell autonomous regulation of retinal ganglion cell and cone photoreceptor fates in the mouse retina. J Neurosci 29, 12865–77.

56. Schick, E., McCaffery, S. D., Keblish, E. E., Thakurdin, C. and Emerson, M. M. (2019). Lineage tracing analysis of cone photoreceptor associated cis-regulatory elements in the developing chicken retina. Sci Rep 9, 9358.

57. Sexton, T., Buhr, E. and Gelder, R. N. V. (2012). Melanopsin and Mechanisms of Non-visual Ocular Photoreception *. Journal of Biological Chemistry 287, 1649–1656.

58. Shimojo, H., Ohtsuka, T. and Kageyama, R. (2008). Oscillations in Notch Signaling Regulate Maintenance of Neural Progenitors. Neuron 58, 52–64.

59. Shimojo, H., Ohtsuka, T. and Kageyama, R. (2011). Dynamic Expression of Notch Signaling Genes in Neural Stem/Progenitor Cells. Frontiers in Neuroscience 5,.

60. Snippert, H. J., Van Der Flier, L. G., Sato, T., Van Es, J. H., Van Den Born, M., Kroon-Veenboer, C., Barker, N., Klein, A. M., Van Rheenen, J., Simons, B. D., et al. (2010). Intestinal Crypt Homeostasis Results from Neutral Competition between Symmetrically Dividing Lgr5 Stem Cells. Cell 143, 134–144.

61. Sulston, J. E., Schierenberg, E., White, J. G. and Thomson, J. N. (1983). The embryonic cell lineage of the nematode *Caenorhabditis elegans*. Developmental Biology 100, 64–119.

62. Takatsuka, K., Hatakeyama, J., Bessho, Y. and Kageyama, R. (2004). Roles of the bHLH gene Hes1 in retinal morphogenesis. Brain Res 1004, 148–55.

63. Turner, D. L., Snyder, E. Y. and Cepko, C. L. (1990). Lineage-independent determination of cell type in the embryonic mouse retina. Neuron 4, 833–845.

64. Wang, M., Du, L., Lee, A. C., Li, Y., Qin, H. and He, J. (2020). Different lineage contexts direct common pro-neural factors to specify distinct retinal cell subtypes. Journal of Cell Biology 219,.

65. Weir, K., Kim, D. W. and Blackshaw, S. (2021). A potential role for somatostatin signaling in regulating retinal neurogenesis. Sci Rep 11, 10962.

66. West, E. R., Lapan, S. W., Lee, C., Kajderowicz, K. M., Li, X. and Cepko, C. L. (2022). Spatiotemporal patterns of neuronal subtype genesis suggest hierarchical development of retinal diversity. Cell Reports 38, 110191.

67. Wu, F., Bard, J. E., Kann, J., Yergeau, D., Sapkota, D., Ge, Y., Hu, Z., Wang, J., Liu, T. and Mu, X. (2021). Single cell transcriptomics reveals lineage trajectory of retinal ganglion cells in wild-type and Atoh7-null retinas. Nature Communications 12,.

68. Xiang, M. (2013). Intrinsic control of mammalian retinogenesis. Cell. Mol. Life Sci. 70, 2519–2532.

69. Yang, Z., Ding, K., Pan, L., Deng, M. and Gan, L. (2003). Math5 determines the competence state of retinal ganglion cell progenitors. Developmental Biology 264, 240–254.

70. Yaron, O., Farhy, C., Marquardt, T., Applebury, M. and Ashery-Padan, R. (2006). Notch1 functions to suppress cone-photoreceptor fate specification in the developing mouse retina. Development 133, 1367–1378.

71. Young, R. W. (1985). Cell differentiation in the retina of the mouse. The Anatomical Record 212, 199– 205.

72. Zhang, Q., Zagozewski, J., Cheng, S., Dixit, R., Zhang, S., De Melo, J., Mu, X., Klein, W. H., Brown, N. L., Wigle, J. T., et al. (2017). Regulation of *Brn*3b by *Dlx*1 and *Dlx*2 is required for retinal ganglion cell differentiation in the vertebrate retina. Development 144, 1698–1711.

73. Zheng, M.-H., Shi, M., Pei, Z., Gao, F., Han, H. and Ding, Y.-Q. (2009). The transcription factor RBP-J is essential for retinal cell differentiation and lamination. Molecular Brain 2, 38.

